# Dorsal prefrontal cortex drives perseverative behavior in mice

**DOI:** 10.1101/2024.05.02.592241

**Authors:** A. Lebedeva, Y. Wang, L. Funnell, B. Terry, Y. J. Oh, K. Miller, K.D. Harris

**Affiliations:** UCL Sainsbury Wellcome Centre; UCL Institute of Neurology; UCL Institute of Ophthalmology; Google DeepMind

**Author notes:** Co-senior authors (K. Miller, K.D. Harris).

## Abstract

Perseveration – repeating one choice when others would generate larger rewards – is a common behavior, but neither its purpose nor neuronal mechanisms are understood. Here we demonstrate a neural correlate and causal role of dorsal prefrontal cortex, specifically anterior supplementary motor cortex (MOs) in perseveration in mice performing a dynamic reward learning task. An auditory go cue signaled mice to turn a wheel either left or right, with the reward probability of each action switching in blocks. Mice perseverated, gaining suboptimal reward, but were faster when making repeated choices. Neuropixels recordings found neurons whose activity correlated with perseveration and predicted rapid reaction times, almost exclusively in anterior MOs. Optogenetically inhibiting this region at choice time reduced perseveration and slowed reactions. In contrast, inactivating medial prefrontal cortex at choice time had no effect, but inactivating it after reward delivery impaired learning. In this task, therefore, anterior MOs reflects a perseverative decision variable, and is necessary for mediating the effect of this decision variable on choice and reaction time.

---

The psychologist Edward Thorndike proposed two laws of learning (Thorndike, 1911). The “law of effect” holds that when reward follows an action, that action becomes more likely in the future. The less well-known “law of exercise” holds that just performing an action, even without reward, increases the probability this action will be performed again.

The law of effect has inspired substantial research in psychology and neuroscience. Much of this work has applied computational models inspired by reinforcement learning (RL; Sutton & Barto, 1998; Botvinick et al., 2020) to predict how reward history shapes choices, and interpret neural activity recorded during learning tasks (Averbeck & O’Doherty, 2022; Costa & Averbeck, 2020; Daw, 2011; Ito & Doya, 2011; Lee et al., 2012). This work has identified correlates of RL-related cognitive variables in frontal cortex (Bari et al., 2019; Barraclough et al., 2004; Behrens et al., 2007; Costa & Averbeck, 2020; Daw et al., 2006; Seo et al., 2007; Sul et al., 2010, 2011), suggesting a key role of frontal cortex in reward-guided learning and decision-making.

The law of exercise has received comparatively less attention. This law predicts “perseverative” learning: the mere act of performing an action, even in the absence of reward, causes that action to become more likely in the future. Behavior consistent with perseverative learning has been reported in many species and tasks (e.g. Aarts et al., 1998; Gold et al., 2008; Lau & Glimcher, 2005; Padoa-Schioppa, 2013; Riefer et al., 2017), and has been quantitatively described by computational models (Ashby et al., 2007; Ashby et al., 2010; Miller et al., 2019; Bogacz, 2020). The benefits of perseveration to an animal are unclear. One possibility is that repeatedly performing an action makes it cognitively simpler or physically easier (Ashby et al., 2007; Gershman, 2020; Wood & Rünger, 2016), consistent with the observations that repeated actions are performed more reliably and quickly (Bertelson, 1965; Gore et al., 2002; Hikosaka et al., 1995), and that responses under time pressure are more likely to be perseverative (Hardwick et al., 2019; Lai & Gershman, 2024).

Relatively little is known about the neural mechanisms of perseveration. A key question is to understand which brain regions carry neural activity that correlates with, and has a causal role in, perseverative learning and decision-making. In particular, it is unclear whether perseveration is supported by a similar set of brain regions and internal cognitive variables as reward-guided learning and decision-making.

Here we address both questions using a dynamic reward learning task (the “probabilistic reversal learning” task; Samejima et al., 2005; Ito & Doya, 2009; Kim et al., 2009) in head-fixed mice. Choices were largely predicted by past choices, with a much smaller influence of past rewards, and could be well approximated by behavior models that incorporated both perseverative and reward-guided learning. Spiking activity in anterior secondary motor cortex (MOs) correlated with perseveration, including during the inter-trial period when no movements were allowed, but such correlations were largely absent from the other frontal regions. Optogenetically inactivating anterior MOs, but not medial prefrontal cortex (mPFC), during choice attenuated both reward-guided behavior and perseveration, and also weakened the perseverative effect of the current choice on future trials. Inactivating MOs following the reward had no discernible effect, but inactivating mPFC following the reward slowed learning, if delivered for multiple trials in succession.

Together, these results indicate that the anterior secondary motor cortex, uniquely among frontal cortical regions, carries a perseveration decision variable that is used to guide choice in our task. Perseverative learning likewise depends on anterior MOs, but reward-guided learning seems to depend instead on other brain areas including mPFC.

## Results

### Choices in a dynamic reward-learning task are driven largely by perseveration

We trained mice in a head-fixed probabilistic reversal learning task (Figure 1). The mouse was head-fixed, with forepaws on a rubber wheel, and a tube near its mouth for delivering liquid rewards (Figure 1A). To initiate each trial, the mouse held the wheel still for a 200ms Fixation Period. An auditory Go Cue signaled the mouse should turn the wheel either clockwise or counterclockwise (from the mouse’s perspective; we call these left and right choices). After turning the wheel a sufficient distance, the mouse received either a liquid reward or auditory white noise indicating that no reward would be delivered. On each trial, one turn direction (Correct) led to 80% reward probability, while the other (Incorrect) led to 20% reward probability (Figure 1B). Which action (left or right) was correct remained stable for a block of trials, then reversed. The first block of each session lasted between 1 and 175 trials, with subsequent blocks lasting between 125 and 175 trials, each sampled independently from a uniform distribution (Figure 1C).

**Figure 1.**
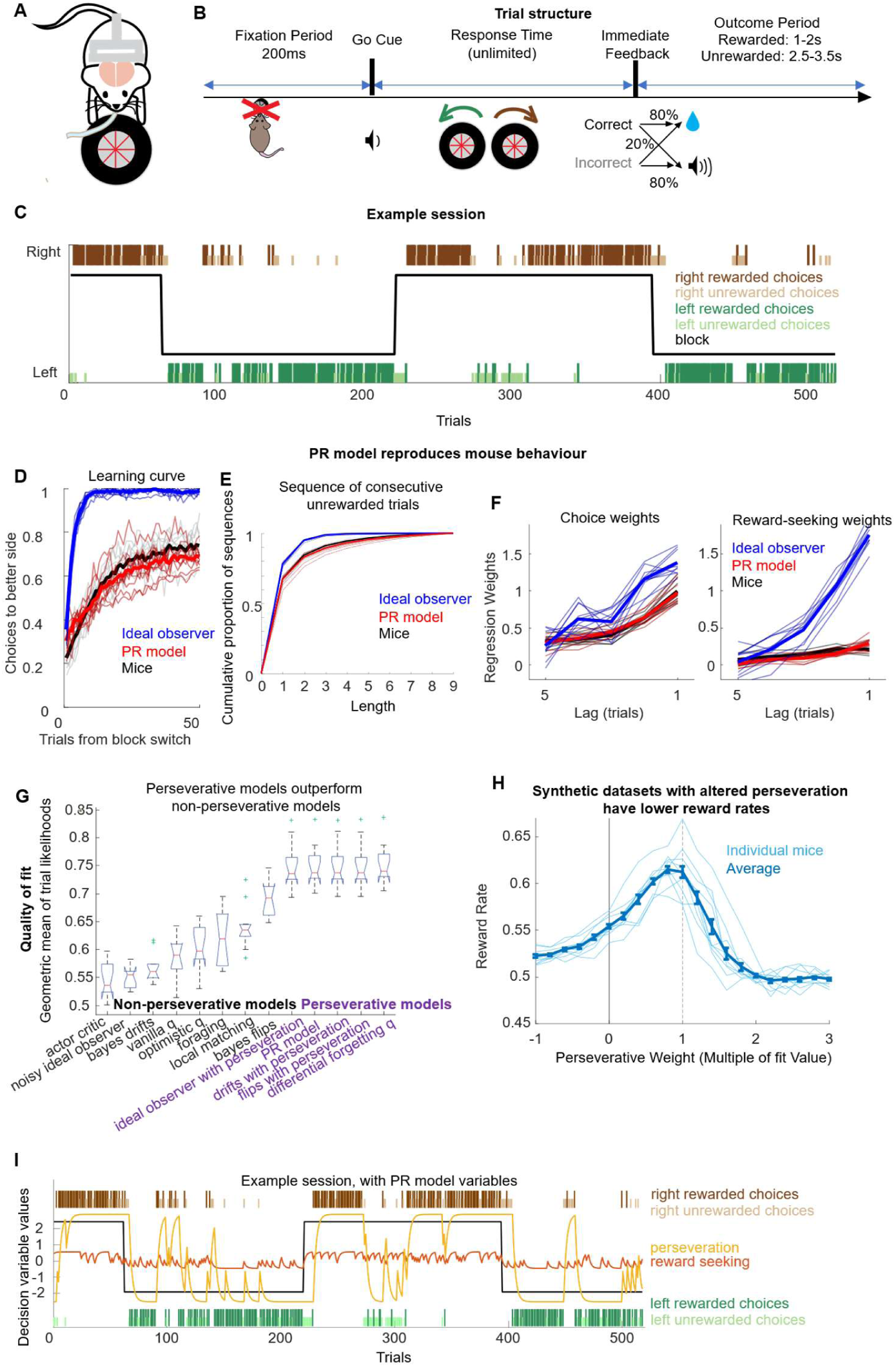
Probabilistic reversal learning task. **A,** The mouse is head-fixed with paws on a wheel and a spout for delivering reward. **B**, To start a trial, mice must hold the wheel still for 200ms. An auditory Go Cue (0.1 s, 5 kHz) indicates trial start but provides no information about which choice will be rewarded. Feedback (reward or white noise) is given after mice turn the wheel 18mm in either direction, with no time limit. Reward probability depends on the choice, switching in blocks. **C**, Example behavioral session. **D,** learning curves aligned to block switches for mice (black), PR model (red) and ideal observer model (blue). **E**, Lengths of unrewarded trial sequences. **F**, Trial-history regression weights, predicting choice on trial N from choice, rewards, and their interactions on trials N-5 to N-1. Left: choice weights; right: interaction weights. Interaction weights of the mouse data were higher that the choice weights for every lag (paired t-test, n=10 mice, p<1e-4). **G**, Quality of fit of 13 probabilistic behavioral models to observed choices sequences, assessed by geometric mean likelihood on held-out trials. Boxes show distribution across sessions. Central mark: median; box edges: 25th and 75th percentiles, green +: outliers, whiskers: range excluding outliers. **H,** Synthetic datasets with altered perseveration have lower reward rates. **I**, Internal state of the PR model fitted to the example session of Fig. 1C.

Mice performed suboptimally in the task. To show this, we compared their choices to those of an ideal observer model, which optimally infers based on history which side is most likely to be Correct, and deterministically chooses this side. The ideal observer earned reward on 77% of trials, substantially outperforming the mice at 62±0.6% (n=10 mice; mean ± S.E.). Following a block switch, mice persisted in choosing the previously-Correct option for much longer than the ideal observer, taking 11.3 ± 0.2 (n=10 mice; mean ± S.E.) vs. 2.2 trials to reach 50% accuracy, and reaching an asymptotic accuracy of 75% ± 0.01% vs 97% correct choices by 100 trials into the block (Figure 1D,E). We characterized the interplay of reward seeking and perseveration using trial-history regression (Lau & Glimcher, 2005; Lee et al., 2004) to predict upcoming choices using the history of previous rewards, choices, and reward-choice interactions (i.e. 1 for a rewarded right choice or unrewarded left choice, −1 for an unrewarded right or rewarded left choice, which indicates the likely correct side). Fitting this model to mouse behavior revealed the choice weights were consistently higher than the reward-choice interaction weights, which was not the case for the ideal observer model (Figure 1F). Overall, these diagnostics reveal that mouse choices in this task show a strong tendency to resemble previous choices, suggesting a role for perseverative learning.

### Mouse choices are well fit by a perseverative behavior model

We quantitatively summarized the interplay of reward-seeking and perseveration by fitting probabilistic trial-by-trial cognitive models (Daw, 2011). Because it was *a priori* unclear which of the many models in the literature would provide the best fit, we evaluated the performance of 13 models using cross-validated likelihood (Figure 1G). The difference in fit of the 13 models was highly significant (p=8×10^-40^, ANOVA), but there was no significant difference in the fits of the five top models, all of which incorporated a perseverative learning mechanism: Perseveration/Reward-learning model (PR model, (Miller et al., 2021)); differential forgetting Q-learning (Ito & Doya, 2009); ideal observer with perseveration (see Methods); ideal-observer Bayesian filters with perseveration for either a generic probabilistic reversal-learning task (see Methods), or a generic drifting reward probability task (Beron et al., 2022). To further evaluate these models, we analyzed synthetic behavioral datasets generated by each model, using parameters fit to each mouse. The five winning models produced similar behavioral sequences to the mice, whether assayed by blockwise learning curves (PR Model: Figure 1D, red curves; remaining models, Supplementary Figure 1), distribution of unrewarded sequence lengths (PR model, Figure 1E; remaining models, Supplementary Figure 1), or trial-history regression weights (PR Model: Figure 1F). In contrast, each of the eight non-winning models showed behavior that deviated in some way from that of the mice (Supplementary Figure 1), falsifying these models (Palminteri et al., 2017). Although the five winning models were not mathematically equivalent, the choice probabilities they predicted were highly correlated (Supplementary Figure 2), suggesting that they captured similar aspects of behavior.

Given that the five perseverative models were equally successful, we chose the simplest among them, the PR model (for Perseveration/Reward learning; Miller et al., 2021). Although this model’s performance was not significantly higher than the other winning models’, it is the simplest model which uses separate decision variables for reward-learning and perseverative components, which will facilitate later analyses of the neural correlates of these. The PR model has two decision variables: *P* which follows the law of exercise and mediates perseverative behavior, and *R* which follows the law of effect and mediates reward-seeking behavior. On each trial, the PR model agent chooses right over left with log odds given by the sum of the two decision variables:

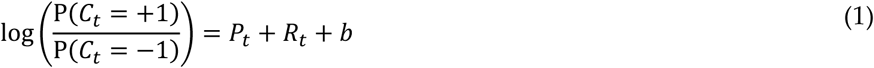

where *b* is a parameter governing fixed left/right bias, *C*_*t*_ is the choice on trial *t* (−1 or +1 for left or right). The perseverative variable *P* is initialized to zero on the first trial, and updated according to:

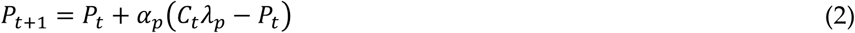

where *⍺*_*p*_ is a learning rate parameter governing the extent to which an individual choice affects *P*, and *λ*_*p*_ is a magnitude parameter governing the largest possible magnitude of *P*. This rule therefore updates *P* towards −*λ*_*p*_ following left choices and +*λ*_*p*_ following right choices, so *P* can be thought of as a running average of previous choices (Figure 1I gold line).

The reward-seeking decision variable R is likewise initialized to zero, and updated according to:

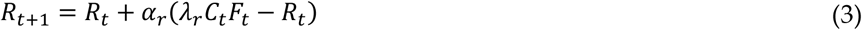

where *F*_*t*_ is the feedback on trial *t* (−1 or +1 for unrewarded or rewarded), *⍺*_*r*_ and *λ*_*r*_ are learning rate and magnitude parameters, which can differ from the perseveration parameters *⍺*_*p*_ and *λ*_*p*_. The rule updates *R* towards −*λ*_*r*_ following rewarded left choices or unrewarded right choices, and towards +*λ*_*r*_ following rewarded right choices or unrewarded left choices. Thus, *R* can be thought of as a running average of the choice-reward interaction, which provides an estimate of the correct choice (Figure 1I, orange line). The overall magnitudes of *P* and *R* – determined by the parameters *λ*_*p*_ and *λ*_*r*_ – therefore govern the relative influence of perseveration and reward seeking on choice.

We fit the model parameters to the observed behavior of each mouse by maximum likelihood, and found that the perseverative process had more than three times the influence of the reward-seeking process (ratio of *λ*_*p*_ to *λ*_*r*_ : 3.67 ± 0.44, mean ± S.E.M., n=10 mice; *λ*_*p*_ greater than *λ*_*r*_ in 10/10 mice; Supplementary Figure 3A), and the learning rate of the perseverative process was somewhat lower than that of the reward-seeking process, meaning that it was influenced by a larger number of past trials (ratio of *⍺*_*r*_ to *⍺*_*p*_ : 1.33 ± 0.13 (mean ± S.E.M., n=10 mice, Supplementary Figure 3A).

Exploring the parameters of the PR model suggested that perseveration increases rather than lowers reward rates, but only because the mice’s reward learning parameters are suboptimal. To show this we simulated behavioral datasets from the PR model with all parameters set to those fit to mouse behavior, except for the perseverative weight *λ*_*p*_, which we swept through a range of values (Figure 1H). We found that both very high and very low values of perseveration decreased reward rate, and that maximum reward rate was achieved using values of the perseverative weight near those used by the mice. Nevertheless, the reward rates obtained by the PR model with mouse-fit parameters (and by the mice themselves) could be exceeded by increasing the reward-seeking weight *λ*_*R*_ and decreasing the reward-seeking learning rate *⍺*_*R*_, following which perseveration offered no further benefit (Supplementary Figure 3B-D). We conclude that the strategy the mice use is suboptimally weighted to perseveration over reward learning, but that given low reward-seeking weights, perseveration helps rather than hinders performance. The number of sessions required to master the task did not correlate with the strength of perseveration of the trained mice (Supplementary Figure 4).

### Perseverative choices are associated with faster responses

Previous work has found that repeated choices are made faster (Bertelson, 1965; Cho et al., 2002; Hikosaka et al., 1995) and that responses made under time pressure are more likely to be perseverative (Hardwick et al., 2019; Lai & Gershman, 2024). To test this in our task, we quantified each trial’s response time as the time between go cue onset and when the wheel turn magnitude passed the threshold required to register a choice (Figure 2A, B). We found that response times were longer on “switch” trials, where the mouse selected a different action than on the previous trial, than on “repeat” trials, where it selected the same action (Figure 2C; switch trials 2.5±0.3 mean±s.e., repeat trials 4.5±0.4 mean±s.e., p = 0.002, signed rank test, n=10 mice). This was not limited to the switch trials themselves: response times remained slow for three trials following the switch and were also slow for trials immediately prior to the switch (Supplementary Figure 5A). We confirmed that when the perseveration variable P had large absolute value, indicating a sequence of choices to the same side, response times were shorter (Pearson correlation −0.15±0.02, mean ± s.e. over n=10 mice; p = 3×10^-4^, t-test; Figure 2D). The absolute reward seeking variable R did not significantly correlate with response time (−0.001±0.02, mean ± s.e. over n=10 mice, p = 0.96 t-test; Figure 2E). Thus, response times in our task are shorter when mice perseverate.

**Figure 2.**
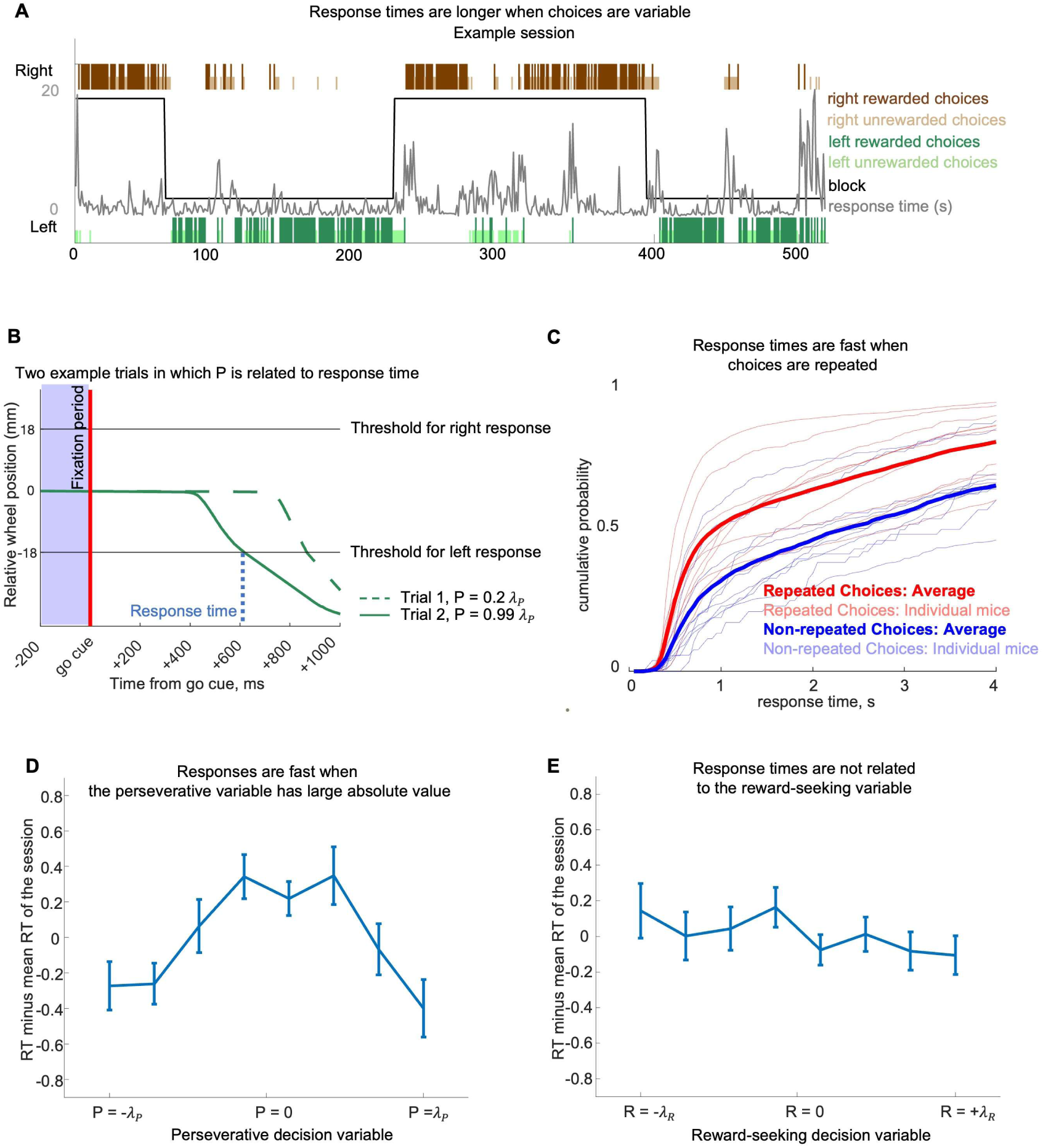
Responses are faster for repeated actions, where the perseverative variable has large magnitude. **A**, Same example session as Fig. 1C, with response times shown in gray. **B**, wheel position on two example trials (dashed: a right turn following LLRLR; solid: a right turn following RRRRR) with different values of the perseverative variable (P). **C**, Cumulative probability density of response times for repeat trials (red) and switch trials (blue), truncated at 4s. Thin lines, individual mice; thick line, average. **D**, Response time for trials with different values of perseveration variable *P* in the PR model minus mean response time (mean ± s.e., n=10 mice). **E**, Response time for trials with different values of reward seeking variable *R* minus mean response time (mean ± s.e., n=10 mice).

### Recordings across the forebrain reveal correlates of future choice specifically in MOs

To search for neural correlates of perseveration and reinforcement learning, we used Neuropixels probes (Jun et al., 2017; Steinmetz et al., 2021) to record across the frontal cortex, as well as several related brain regions, while mice performed the task (Figure 3). After automatic spike sorting (Pachitariu et al., 2016; Pachitariu et al., 2023) and manual curation, our dataset comprised 19,893 units from 42 sessions in 10 mice. We obtained a sufficient number of units and sessions for detailed analysis in five target major brain regions that have been implicated in prior work in decision-making on reward-guided tasks: secondary motor cortex (MOs, 2,580 units in 15 sessions; our MOs recordings targeted an anterior subregion that has been implicated in similar wheel-turning tasks (International Brain Lab et al., 2023; Steinmetz et al., 2019; Zatka-Haas et al., 2021); for brevity we use the unqualified term “MOs” to refer specifically to this subregion throughout the paper), medial prefrontal cortex (mPFC, 4,255 units in 14 sessions; combining prelimbic, infralimbic, and anterior cingulate cortex), orbital cortex (ORB, 2,400 units in 11 sessions; combining medial, ventrolateral, and lateral orbital cortex), dorsomedial striatum (DMS, 1,287 units in 9 sessions), and dorsal hippocampus (DH, 2,176 units in 15 sessions; combining CA1, CA3, and dentate gyrus). We also obtained a sufficient number of recordings from olfactory areas (OLF, 2,509 units in 15 sessions; combining olfactory bulb, olfactory nucleus, taenia tecta, dorsal peduncular area, piriform area, nucleus of the lateral olfactory tract, cortical amygdalar area, piriform-amygdalar area and postpiriform transition area) which lie ventral to many of our target regions (Figure 3B). Other regions recorded include lateral septal nucleus (646 units in 6 sessions), retrosplenial area (185 units in 5 sessions) and nucleus accumbens (429 units in 6 sessions) (Supplementary Figure 6), but the smaller number of sessions and units for these latter areas precluded rigorous statistical analysis.

**Figure 3.**
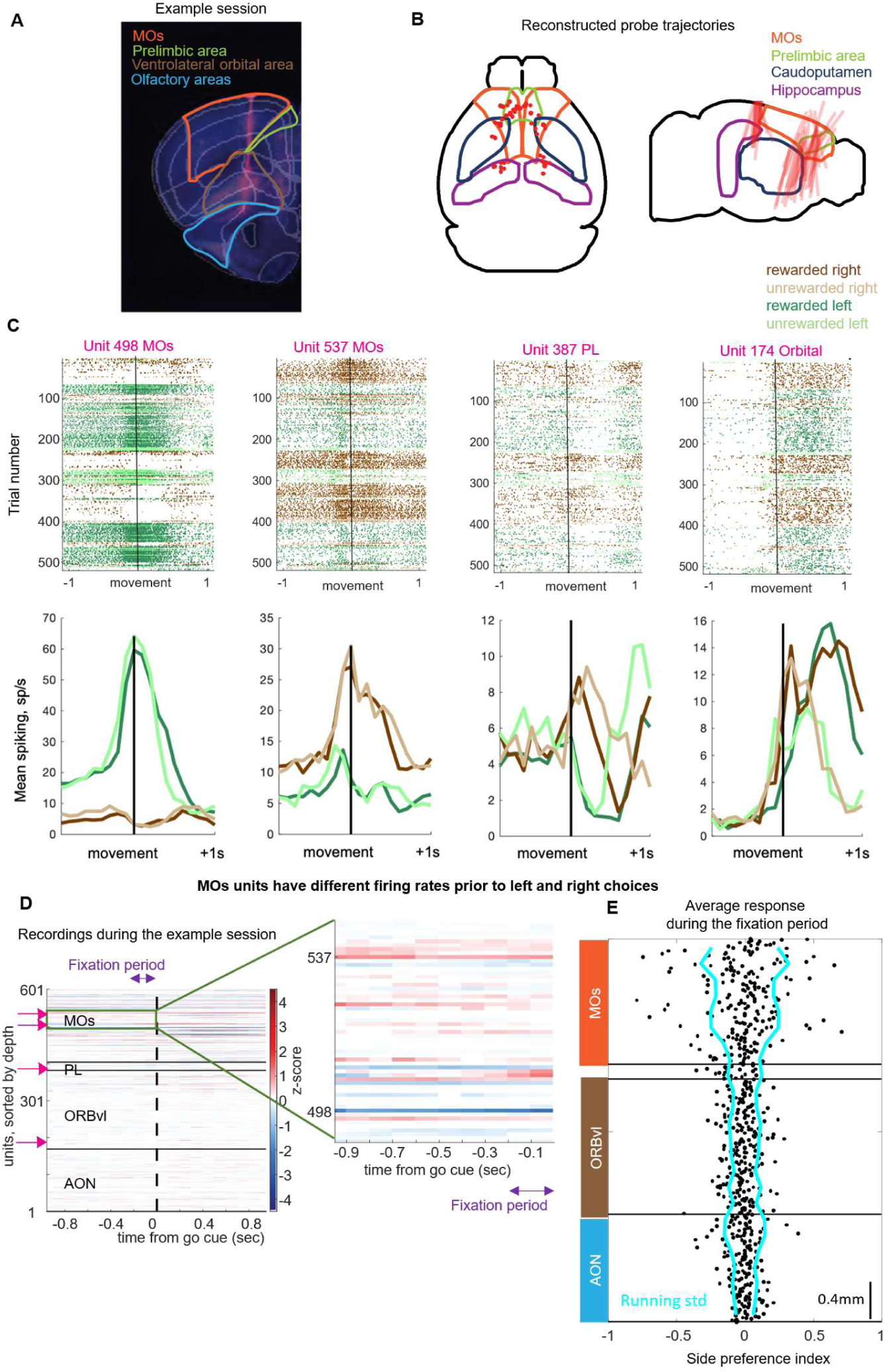
Example MOs neurons showing cue-period correlates of upcoming choices. **A**, Histological electrode track reconstruction for one example mouse. **B**, Reconstructed probe trajectories for all recordings. **C**, Top: raster plots for four example units from the probe in A, trials ordered chronologically and colored by trial type. Bottom: peristimulus time histograms for the same units. Activity of each trial is aligned to the first movement detected on that trial. Time shown in seconds relative to the movement onset. **D,** Difference in z-scored activity prior to right and left choices in an example session, for all neurons as a function of depth. Each time bin was z-scored independently by subtracting the mean and dividing by the standard deviation across depths. Pink arrows (left): units from C. Activity of all units is shown in seconds relative to the time of the go cue on all trials. **E,** Neural preference for future choice side vs. depth. X-axis: side preference index 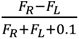, where *F_*R*_* and *F_*L*_* are mean spikes count in fixation periods prior to right and left choices. Cyan curve: running standard deviation (25 units) of side preference as a function of depth.

Consistent with previous work (Lee et al., 2012; Steinmetz et al., 2019), we found example units in all brain regions whose spike rates (defined by peristimulus time histograms) correlated with choice and feedback after these events had begun (Figure 3, Supplementary figures 7, 8, 9). However, example units whose spike rates during the fixation period correlated with upcoming choice were rarer and appeared far more common in anterior MOs than in any other region (Figure 3C-E, Supplementary Figure 10). A similar distinction has been observed previously in rats performing a related freely-moving task (Sul et al., 2011; Lee et al., 2012) and interpreted as indicating a specific role for MOs in value-guided action selection.

Verifying the statistical significance of a correlation between neural firing and choices requires care. Both behavioral and neural timeseries show temporal autocorrelations, including due to artifacts such as electrode drift. Commonly used statistical tests assuming that observations are statistically independent of one another can produce spuriously significant “nonsense correlations” in this situation, even in the absence of genuine correlations (Yule, 1926; Harris, 2021b; Elber-Dorozko & Loewenstein, 2018). For example, consider a unit whose rate was artifactually higher at the start of the experiment due to electrode drift, in a session that started with left choice block. A t-test would show significantly higher rates in left than right choice trials, even if the unit carried no genuine information about the choice. Furthermore, because simultaneously recorded units are correlated, tests which assume independence of units may also yield false positives. We solved this problem using a “session permutation” test (Harris, 2021b), which treats an entire recorded session as the unit of independent variability. We predict the sequence of choices in each session from the sequence of neural population activity vectors (defined as the spike counts of all neurons in a 200ms time window) using logistic regression, and compute a single test statistic summing the log likelihood of all sessions. We compare this statistic to a null distribution obtained by rerunning after permuting the session labels of the behavioral data. This therefore tests the null hypothesis that neural activity in a particular session was no more related to the choices made by the mouse in that session than to choices made in an entirely different session, without assuming independence between trials or units within a session.

This analysis confirmed that fixation-period population activity in MOs predicted the upcoming choice. We applied this analysis to neural activity in each of our brain regions, both during the fixation period before movement had begun, and in 200ms timebins following movement initiation. Once the mouse had begun to turn the wheel, the direction of choice was robustly reflected in neural activity in all of our brain regions (Figure 4a); similarly, all regions correlated with the feedback variable after feedback was delivered (Supplementary Figure 7). However during the fixation period, the direction of upcoming choice was significantly decoded only from MOs (p<1e-5, all other p>0.01; Figure 4A). This did not result from different numbers of units recorded in the different regions: repeating the analysis using an equal number of units from each region still yielded better prediction from MOs than all others (Figure 4B, Supplementary Figure 11). Thus, neural activity in MOs bears a relationship to upcoming choice that is unique among the brain regions we recorded, suggesting that it may play a distinct role in the choice process. We note that the MOs was also the only area in which the upcoming choice, the P variable, or the decision variable P+R could be predicted in the 200ms following the go cue, while 200ms prior to detected movement the choice could be decoded from the activity of all recorded areas (Supplementary Figure 12, 13, 14; recall that we use “MOs” to refer specifically to its anterior aspect, Supplementary Figures 8,9). We also repeated the analysis using Lasso regression and again found that MOs was still the only area where the P variable could be decoded prior to the go cue (Supplementary Figure 15).

**Figure 4.**
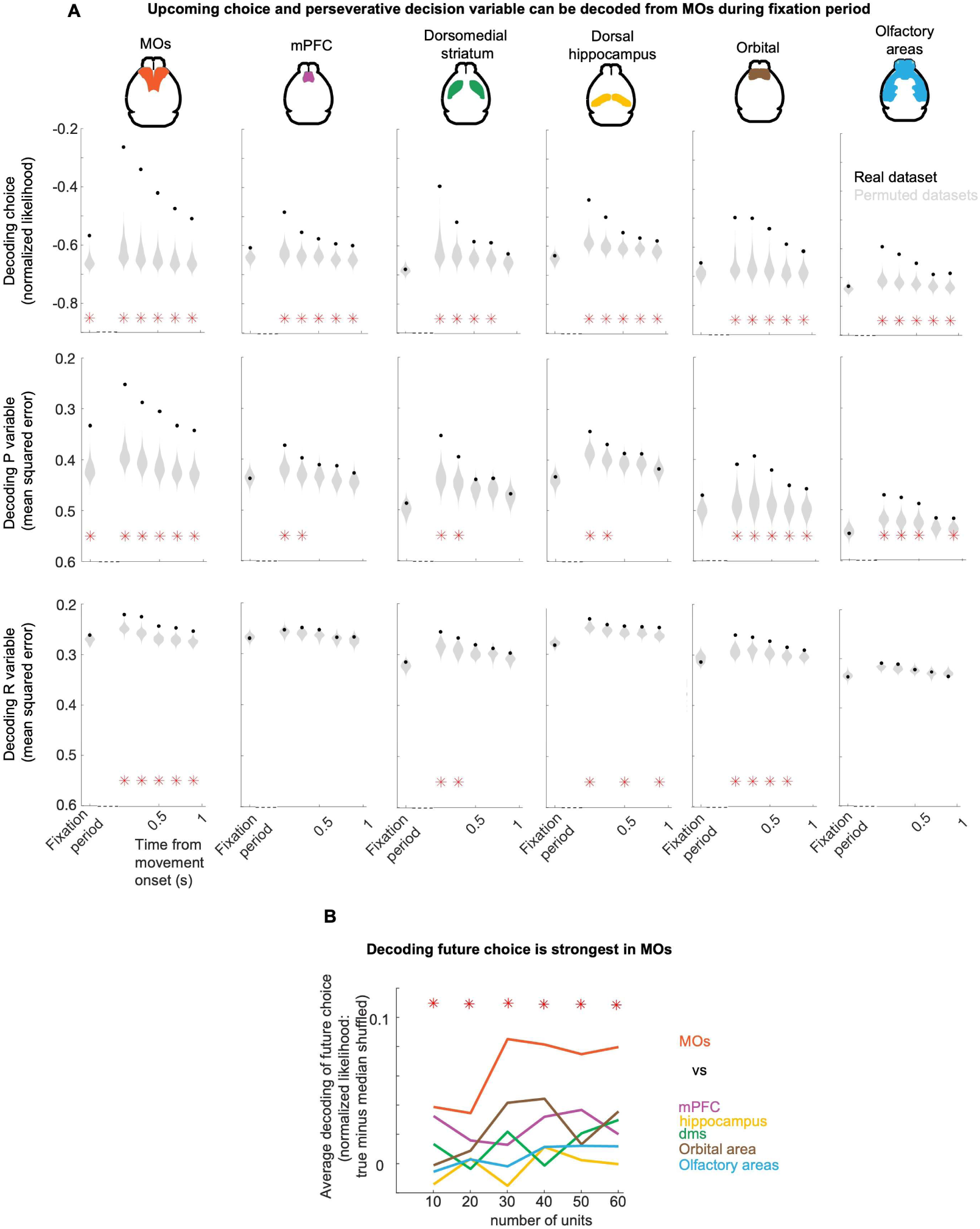
Upcoming movements are encoded specifically in MOs. **A,** Decoding of upcoming choice and model variables from population activity. Each column shows decoding of choice (logistic regression), perseveration variable *P* (linear regression); and reward-seeking variable *R* (linear regression). X-axis: time epoch; fixation period (0-200 ms before Go Cue), and subsequent 200 ms epochs from movement onset. Black dots: mean prediction of actual value across sessions; gray violins, null distribution from session permutation. Red stars: p<0.01, session permutation test. **B,** Decoding upcoming choice from fixation-period activity of equally-sized random subsamples of neurons from each region. X-axis: number of units in subsample; Y-axis, mean log-likelihood relative to permuted sessions. Red stars: MOs significantly different to other regions (p<.01; t-test).

In the PR model, the choice on trial *t* is driven by a weighted sum of the perseverative variable *P*_*t*_ and the reward-seeking variable *R*_*t*_, and patterns of neural firing in MOs during the pre-choice fixation period correlated significantly with *P*_*t*_ but not with *R*_*t*_. To show this we used a linear regression model to relate each session’s sequence of *P* or *R* values to the sequence of neural population activity vectors on the corresponding trials, and summarized this relationship by averaging the mean squared error across sessions. We compared this statistic to a null distribution obtained by permuting the session labels of the behavioral data. After the mouse had begun to turn the wheel, *P* was robustly reflected in neural activity of each brain region (Figure 4A, second row). However, MOs was the only region whose activity robustly reflected *P* during the fixation; (p<1e-5, all others p>0.01). There was no side bias in these correlates: neurons in the left and right hemispheres were equally likely to correlate with perseveration in a leftward or rightward direction (Supplementary Figure 16).

The reward-learning variable *R* was not robustly reflected in any brain region during the fixation period. It was robustly reflected during the movement only in MOs and ORB (Figure 4A, third row). We additionally used the fact that rewards were randomly delivered to test for a causal effect of rewards on subsequent neural activity. This found that reward had a significant causal effect on activity in all recorded regions, that persisted even through the subsequent trial (Supplementary Figure 17). However in MOs, the interaction of choice and feedback – which updates the reward-seeking decision variable *R* – had a more transient causal effect that was no longer significant by the time of the subsequent fixation period (Supplementary Figure 18). We conclude that MOs activity in the fixation period robustly reflects the perseverative decision variable *P*, but has little or no relationship with the reward-seeking decision variable *R*. This indicates that MOs tracks recent actions, consistent with a role in value-free or habitual behavior, but that it does not seem to track recent action-outcome associations, casting doubt on the idea that it plays a role in value-based or goal-directed behavior.

### Secondary motor cortex activity reflects a perseverative decision variable

The fact that the activity of units in MOs correlates with both upcoming choice and the *P* variable, which are themselves correlated, has several possible interpretations. First, MOs might simply reflect the upcoming choice in a binary way, reflecting an internal decision made prior to the fixation period that is awaiting the go cue to be released into motor action; since *P* is correlated with upcoming choice, such a motor intention would correlate with *P* as well. Alternatively, MOs might reflect a continuous perseverative decision variable which itself is correlated with binary future choice.

Examining the activity of individual MOs units revealed cells whose activity varied with *P* beyond their dependence on upcoming choice (Figure 5). An example MOs unit was more active on average prior to left than to right choices but showed further dependence on choice history (Figure 5A). This unit fired essentially no fixation-period spikes after a long sequence of right choices, but it was active following a sequence of left choices, even if the upcoming choice was to the right. Similarly, the mean activity of neurons predicting either left or right choices showed a correlation with *P* even among trials with the same upcoming choice (Figure 5B).

**Figure 5.**
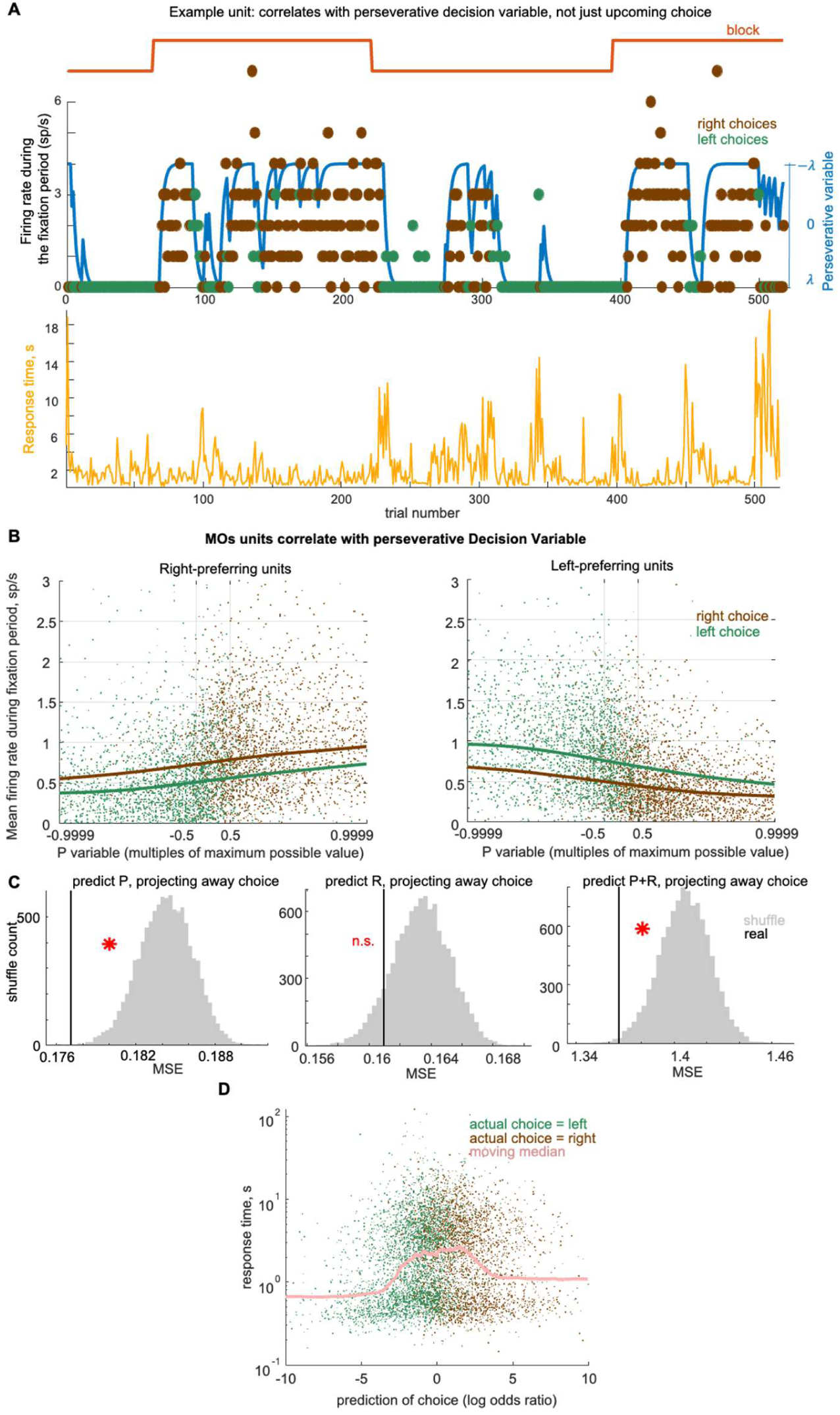
Choice correlates in MOs reflect perseverative decision variable. **A,** Top, example unit whose fixation-period activity correlates with *P*. Bottom, response times in the same session. **B,** Mean activity of left- or right-preferring units during fixation period, as a function of *P* (x-axis; logistic scale) and choice direction (color). Each point corresponds to a trial, curves to a lowess smoothing of the points. **C**, Mean square error (MSE) of predicting *P, R,* or *P+R* after projecting out choice. Black line: actual value; gray histogram, null distribution from session permutation. Stars: p<.01, session permutation test for partial correlation. **D,** Response time as a function of neural choice direction prediction, obtained by logistic regression of choice direction from fixation-period population activity. Brown and green points: single trials with right or left choices; pink curve: running median.

To establish the statistical significance of these results, we used a variant of the session permutation test, designed to identify partial correlations in timeseries with correlation between trials and neurons (Harris & Yuan, 2024). This showed that there was a significant correlation between patterns of activity in MOs during the fixation period and the perseverative decision variable *P,* even after the common effect of upcoming choice was accounted for (Figure 5C). Applying this same analysis to the reward-seeking decision variable *R* did not reveal a significant relationship, consistent with our earlier result that R could not be predicted during the fixation period (Figure 4A). We also asked whether MOs activity could also predict the summed decision variable *P*+*R* when controlling for upcoming choice, and found that it could (Figure 5C, right). These results are therefore consistent with either the hypothesis that MOs fixation-period activity reflects the perseverative decision variable *P* specifically, or that it reflects the aggregate summed decision variable *P* + *R*. Since this aggregate variable is dominated by *P* (Figure 1I, Supplementary Figure 3), both hypotheses indicate that MOs activity strongly correlates with a perseverative decision variable, and correlates with the reward-seeking variable weakly, if at all.

We hypothesized that when MOs activity during the fixation period strongly predicts a given action, such as after a long sequence of choices in the same direction, this should indicate a degree of confidence or preparation of the action, leading to shorter response times. This was supported by an example neuron that appeared to fire strongly during the fixation period prior to trials with short response times (Figure 5A), and also by the observation that both the neural activity in MOs and response time change on trials preceding the choice switch (Supplementary Figure 5A, B). To test the hypothesis statistically, we predicted the mouse’s upcoming choice from fixation-period MOs activity during the fixation period using logistic regression. We reasoned that the weighted sum of activity found by this classifier (prior to passing through the logistic function) would form an estimate of confidence on each trial: when many MOs units are “voting” for a right or left choice, this predictor would have a large absolute value (positive or negative). Even though the predictor was trained without reference to response times, responses were faster on trials when the predictor had large magnitude. (Figure 5D; p=0.02, linear mixed effects model with fixed effect of absolute value of choice prediction and random effects of mouse and session within mouse, n=9 mice). Thus, when pre-cue MOs activity is of the form that strongly indicates either a left or right turn, the motion is executed rapidly, as would be the case if this MOs activity subserved motor preparation for an action to be released as soon as the go cue sounds.

### Inhibition of MOs activity reduces perseveration and slows responses

To investigate the causal role of MOs in the task, we used a transgenic mouse line (Ai32xPV-Cre) which expressed Channelrhodopsin 2 in Parvalbumin-positive interneurons (Guo et al., 2014; Olsen et al., 2012). We implanted six mice bilaterally with cannulas allowing light delivery to the surface of MOs, targeting the same subregion where we recorded correlated neurons; a further six mice had cannulas in the same AP and ML location, but 1mm deeper, to study the effects of medial prefrontal cortex (mPFC) inactivation, and also to act as a control for inactivation spreading to regions immediately beneath the targeted region (Li et al., 2019). After surgery, the mice were trained to perform the task. We defined two behavioral windows for optogenetic inhibition: the “choice period”, from go cue onset until a choice was registered, and the “outcome period”, for 1.6 seconds from the time of feedback (Figure 6B). We applied 473 nm light to suppress neural activity during the choice period on a randomly-selected 20% of trials, and during the outcome period on an independently-selected 20% of trials. Our dataset comprised 44,243 trials from 82 sessions in MOs, and 48,614 trials from 92 sessions in mPFC. Doing inactivations in these two windows within the same session allows us to make direct comparisons between their effects.

**Figure 6.**
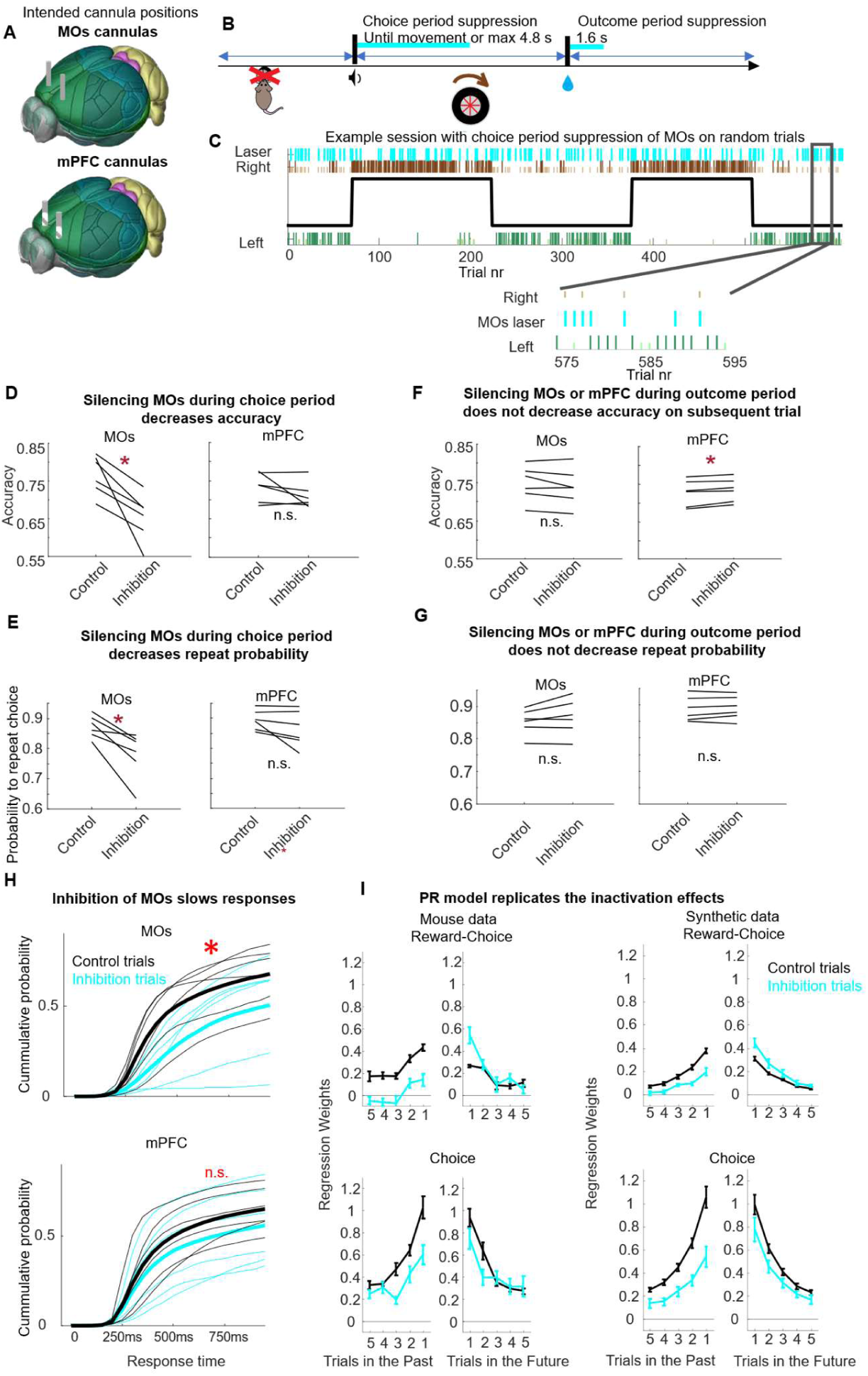
MOs activity is required for perseveration and rapid responses. **A**, Location of light-delivery cannulas in two groups of mice. **B**, Timing of optogenetic suppression in the choice period or outcome period; light was applied independently with 20% probability to each period on each trial. **C**, Example session with choice period-suppression in MOs. Cyan: laser; inset: expanded timescale. **D-E**, Effect of MOs (left) or mPFC (right) suppressing during the choice period on accuracy, i.e fraction of trials choosing the Correct side (D) or probability of repeating the previous choice (E). **F-G**, Same for suppression during the outcome period. **H**, Cumulative histogram of response times on trials with optogenetic suppression in the choice period (cyan) and on no-laser trials (black), for mice with cannulas in MOs (top) and mPFC (bottom). Individual mice, thin lines, average across mice, thick lines. **I**, Left 4 panels, Trial history regression weights for predicting mouse choice using past choices (choice weights), and choice-reward interaction (reward-choice weights), for laser (cyan) and no laser (black) conditions. Left column shows weights from trials N-n to trial N, where inactivation was delivered to MOs on trial N; right column shows weights from trial N to trial N+n, where inactivation was delivered to MOs on trial N. Right 4 panels show the same but for PR model.

Inactivating MOs during the choice period reduced both accuracy of the mice’s choices, and the probability of repeating the previous choice (accuracy in MOs paired t-test, n=6 mice, p = 0.01; repeat probability in MOs paired t-test, n=6 mice, p = 0.01; Figure 6 D-E). Consistent with this observation, the probability to change choice after an unsuccessful trial (“lose-switch” behaviour) was increased upon MOs inactivation (MOs paired t-test, n=6 mice, p= 0.0026; Supplementary Figure 19). In contrast, no significant effect was seen when mPFC was suppressed during the choice period (accuracy in mPFC paired t-test, n=6 mice, p =0.2; repeat probability in mPFC paired t-test, n=6 mice, p =0.12; Figure 6 D-E; Supplementary Figure 19), and the effect of MOs suppression was significantly stronger than that of mPFC inactivation (unpaired t-test comparing mean change in accuracy, n=12 mice, p=0.01). Furthermore, inactivation during the outcome period on random trials, either in MOs or mPFC, did not affect repeat probability, and did not affect accuracy in MOs (accuracy in MOs paired t-test, n=6 mice, p = 0.12; repeat probability in MOs paired t-test, n=6 mice, p = 0.11; repeat probability in mPFC paired t-test, n=6 mice, p = 0.43; Figure 6 F, G). Inactivation of mPFC during the outcome period did not result in a decrease in accuracy, but instead in a very slight increase (average increase 0.65 percentage points, p=0.005; paired t-test, n=6 mice, Figure 6 F, right). The magnitude of accuracy change following mPFC stimulation during the outcome period was significantly smaller than the magnitude of accuracy change during the choice period in MOs (unpaired t-test, n = 12 mice, p = 0.003). Applying trial history regression specifically to trials where MOs was inactivated yielded lower choice weights (expected to be related to perseveration) and lower reward-choice interaction weights (expected to be related to reward-seeking) (Figure 6 I, left panels; choice weights: p=0.01, reward-choice interaction weights: p=0.007, t-test, n=6 mice), but this was not seen with mPFC inactivation (choice weights: p=0.41, reward-choice interaction weights: p=0.79, t-test, n=6 mice). In contrast, the trial history regression applied to trials with and without mPFC inactivation did not show significant differences in either choice or interaction weights (Supplementary Figure 20). Thus, of the inactivation protocols we tested, accuracy and repetition were decreased specifically by inactivation of MOs during the choice period. Inactivation of MOs during the outcome period serves as an internal control, showing that the timing of inactivation mattered and that simply inactivating the region outside the decision window did not affect the choice. Inactivation of mPFC provides a second internal control, showing that the light itself or laser shutter sound was not sufficient to drive change in animal behaviour.

MOs inactivation during the choice period also decreased future perseveration: inactivating MOs on a trial reduced the effect of that trial’s choice on future choices. To show this, we refit the trial history regression, with different weights from inactivated and non-inactivated trials onto future choices. The choice weight from an inactivated trial to the next trial’s choice was decreased, while the reward-choice weight to the next trial’s choice was increased (Figure 6I, right panels; choice weights: p=0.008, reward-choice weights: p=0.01, t-test, n=6 mice). Thus, inactivating MOs during the choice period of a given trial reduces perseverative learning, making the mouse less likely to perseverate and more likely to be reward-driven in the future based on the choice it made on that inactivated trial.

Inactivating MOs during the choice period also caused an increase in response times (Figure 6H). MOs inactivation did not prevent movement: on 84% of trials, the mice turned the wheel while the laser was still on. However, the time at which the wheel was turned was later (2.58±0.58 vs 1.91±0.15 s; median ± s.e., p=0.03, signed rank test, n=6 mice). On a further 16% of laser trials, the mice only made their choices after the laser was switched off (4.8 s after the go cue); the fraction of correct choices was similar on these trials as for the trials with responses during laser stimulation (0.66±0.02 vs 0.74±0.03, p=0.21, signrank test, n=6 mice). When mPFC was inactivated, we observed a trend toward shortened response time although it was not significant (2.96±0.45 vs 2.31±0.25 s, median ± s.e., p= 0.06, signrank test). Together, these results indicate that neural activity in MOs plays a role in guiding and rapidly executing choices, based on both perseverative and reward-seeking strategies; and that MOs activity during the choice has an impact on future choices but only for the perseverative component.

Inactivation following feedback delivery showed that mPFC, but not MOs is involved in reward-based learning. Inactivation of either region during the outcome period (1.6 s following delivery of reward or white noise feedback) on random trials did not have a large effect on subsequent choices (Figure 6F). However, because it takes mice several trials to learn about a block switch, an effect on reward learning might only be observable when multiple trials were inactivated successively. We therefore performed new experiments silencing outcome-period activity on all trials of a randomly selected half of each session (Figure 7A-B). This allowed us to compare learning curves for block switches within the inactivated and non-inactivated half-sessions (Figure 7C). Inactivating mPFC significantly slowed learning about the block switch, but inactivating MOs had at most a marginal effect (Figure 7D), and the difference between the effects of mPFC and MOs inactivation were significant (Figure 7E, p=0.04, t-test, n=6 mice in MOs group, n=7 mice in mPFC group). Thus, while MOs appears to be involved in perseveration and motor execution, mPFC seems to be required for reward learning in this task.

**Figure 7.**
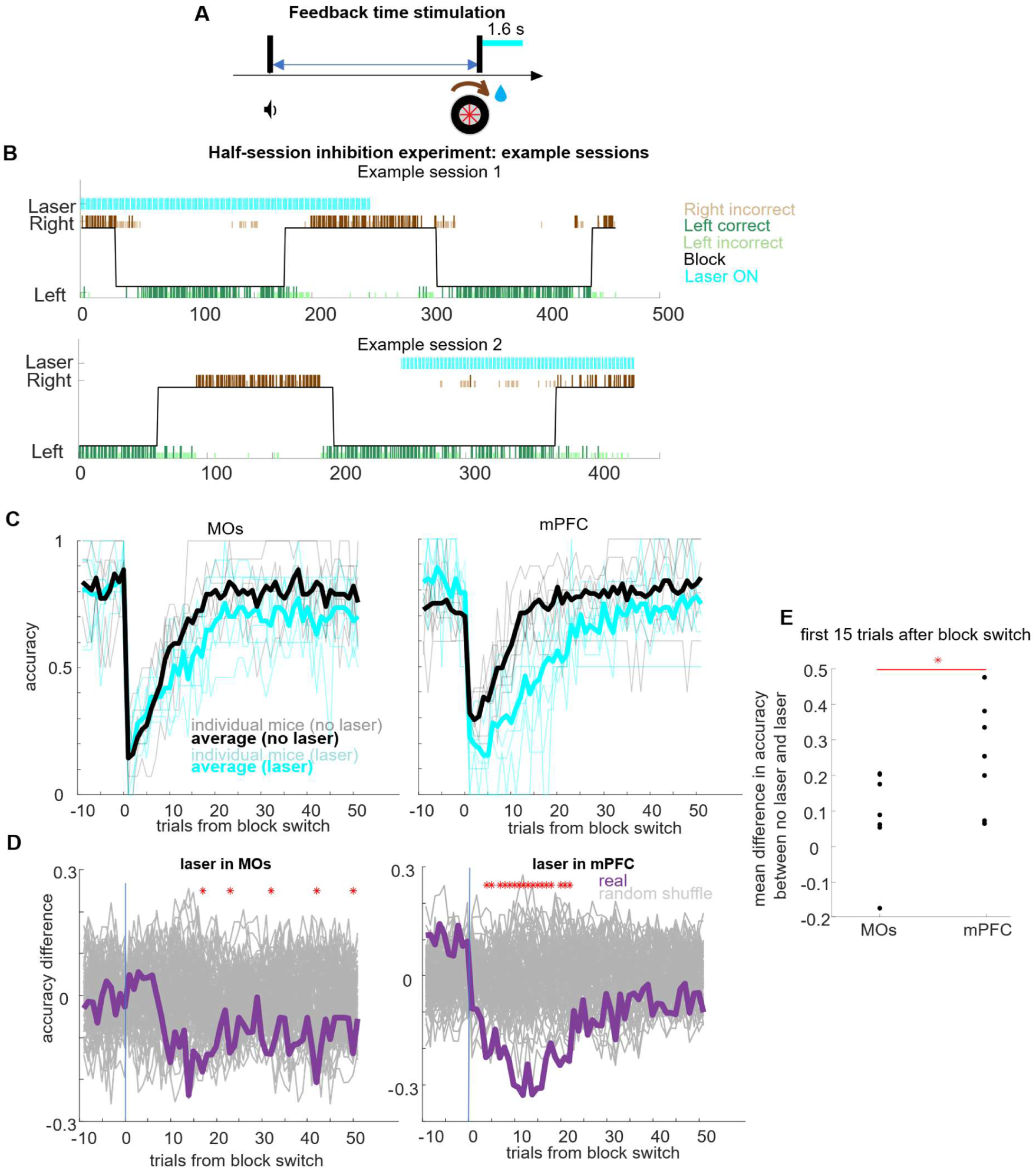
Optogenetic mPFC suppression during outcome period increase the number of trials needed to reach good performance after block flip. **A,** Timing of laser pulses in this experiment. **B,** Two example sessions. For each session, light is applied randomly on either every trial in the first half session (top) or every trial in the second half (bottom). Cyan lines: laser trials. **C,** Learning curve of MOs (left) and mPFC (right) groups, for no-laser (black) and laser blocks (cyan). Thin lines, individual mice, thick lines, average. **D,** Difference in accuracy between laser and no laser blocks averaged over sessions (magenta), compared to a null distribution obtained by randomly resampling which half of trials was inactivated on each session (grey). Left: MOs suppression, right: mPFC suppression. Red Stars: timepoints for which accuracy in the inactivated half-session was worse than the null distribution (p<.01, randomization test). **E,** comparison of accuracy with MOs and mPFC inactivation. Each point shows the difference in accuracy between laser and non-laser halves, averaged over the first 15 trials following each block switch for all sessions of one mouse. The difference between mPFC and MOs was significant (p<.05; rank sum test, n=13 mice).

## Discussion

We found that mouse behavior in a probabilistic reversal learning task could only be captured by computational models containing both reward seeking and perseverative mechanisms. Response times were not correlated with model-predicted reward-seeking drive but were shorter for repeated choices and when the model predicted strong perseverative drive. Neuronal activity in many brain regions correlated with choice once movement had begun, but correlates prior to movement onset were primarily found in the anterior aspect of dorsal prefrontal cortex (MOs). This pre-movement activity was correlated with an analog perseverative decision variable, even when controlling for the binary upcoming choice, but was not correlated with choice when controlling for the decision variable. Silencing MOs, but not mPFC activity at choice time slowed response times and made choice direction closer to random. Silencing mPFC, but not MOs following reward delivery slowed learning rates. Together, these results indicate that MOs is involved in driving perseverative behavior in this task.

The benefits of perseveration to an animal are unclear. It is widely held that perseveration occurs because strategy-switching incurs a cost, either physical (for example because body posture and motor preparation is more effective for repeated actions), or cognitive. The amount of perseveration animals exhibit varies between species, tasks and contexts, and in the current study we observed stronger perseveration than some previous studies of reversal learning. For example, rats tended to alternate rather than perseverate in a real-space probabilistic reversal learning task (Sul et al., 2011), which may be due to rodents’ general tendency to alternate on T-mazes (Deacon & Rawlins, 2006; Dember & Richman, 1989). We also observed stronger perseveration than some other studies using head-fixed reversal learning tasks (Bari et al., 2019; Beron et al., 2022; Hattori et al., 2019; Le et al., 2023). This might be related to our use of longer blocks, or perhaps the steering wheel apparatus might impose a larger cost of switching between choices than left/right licking. The question of whether perseveration is adaptive or maladaptive in the current task is somewhat complex. The mice’s strategy led to less reward than would have been obtained if they had followed the PR model with a higher reward-seeking and lower perseverative weight. Nevertheless, given the too-low reward-seeking weight we actually observed, perseveration did provide an improvement over a reward-seeking-only strategy. Consistent with this, optogenetic inactivation of MOs reduced reward rates as well as reducing perseveration. It is thus possible that perseveration serves to compensate for a suboptimal underestimation of environmental variability in the current task. Regardless of the reason, the strong perseveration observed in this task offered an opportunity to study its neural substrates, which strongly implicated anterior MOs. As discussed below, however, we hypothesize that this arises from a flexible role for MOs in implementing context-appropriate decision rules, which are perseverative in this task but not necessarily others.

Several models for the neural mechanisms of perseveration have been proposed. Some suggest that perseverative information is stored in the strengths of either cortico-cortical (Ashby et al., 2007, 2010) or cortico-striatal synapses (Bogacz, 2020). These models do not require spiking correlates of perseverative information in intertrial intervals. We did however find such correlates in MOs, even before the go cue. This does not preclude a change in synaptic weights, but does raise the possibility that short-timescale perseveration might rely at least partially on persistent activity (J. X. Wang et al., 2018). Recent experiments have shown tail-striatal dopamine correlates with action prediction error, which reflects the difference between the P variable and the action made (Greenstreet et al., 2022). It is possible that MOs correlates of the P variable contribute to the computation of this dopamine signal.

Although many computational models of perseveration propose that it results from the Law of Exercise, whereby performing an action causes that action to become more likely in the future (Ashby et al., 2007; Ashby et al., 2010; Miller et al., 2019; Bogacz, 2020), an alternative posits that animals exhibit discrete behavioral states: “explore” states in which they sample all available options with high probability and “exploit” states in which they sample predominantly from a single option (Ebitz et al., 2018; Zid et al., 2024). In this interpretation, perseveration results from having a low probability to transition from an exploit state back to the explore state. This interpretation is compatible with the types of behavior that would also be seen with Law of Exercise learning, but only if the likelihood to remain in the exploit state increases over time. Our analysis of mouse behavior is thus broadly consistent with either interpretation. However our finding that MOs activity encoded not just the upcoming binary choice but the continuous *P* variable which grows with the number of previous choices in one direction, suggests that any explore/exploit mechanism would need to incorporate a continuous measure of certainty for exploitation, at which point differences to a Law of Effect model become less clear.

Our inactivation results are opposite to those of Sul et al., 2011, who found that in a spatial reversal learning task, MOs lesions increased, rather than decreased, rats’ tendency to perseverate; similarly they observed that MOs lesion decreased lose-switch behavior while we saw inactivation increased it. Although this might reflect a difference in species or inactivation method (static lesion vs. trial-specific optogenetics), we hypothesize that it relates to a difference behavior induced by the experimental apparatus. Specifically, their use of a spatial T-maze and a baiting-based reward scheme both encourage choice alternation rather than consistent selection of a higher-reward option. We speculate that in such anti-perseverative tasks, MOs might still encode the perseverative variable *P*, but that this variable would be negatively weighted by decision circuits. Although Sul et al did not explicitly test this hypothesis, we note that they observed stronger correlates of prior choice than of prior reward or reward-choice interaction, which is consistent with the idea that MOs reflects a perseverative decision variable but not a reward-seeking one. Taken together, these results therefore suggest that MOs might be involved in both perseverative and anti-perseverative strategies, depending on the task, species, and context.

Our neural recording results also contrast with those of Bari et al. (2019), who found that neural activity in mPFC correlates with a decision variable strongly and stably throughout the inter-trial interval, while neural activity in a subregion of MOs correlates much less strongly. In contrast, we find correlates of the decision variable (P+R and/or P) are strongly present in MOs during the analogous period but are largely absent from mPFC during this time. We hypothesize that this again reflects a difference in tasks: Bari et al.’s task involved a “baiting” scheme similar to Sul et al. (2011), and mice showed little perseveration but instead a tendency to avoid the choice that was taken on the immediately previous trial. The observation that MOs does not correlate with decision variable in this task, while it does in ours, is consistent with the idea that this decision variable may be specific to perseverative behavior.

A prominent theory holds that MOs is responsible for short-term memory of information relevant for guiding choice (Barthas & Kwan, 2017; Yang & Kwan, 2021), which is supported by recordings in many tasks (Barthas & Kwan, 2017; Ebitz et al., 2018; Erlich et al., 2011; Li et al., 2015; Murakami et al., 2014; Sul et al., 2011). Our finding that fixation-period MOs activity correlates with the perseverative variable *P* is broadly consistent with this idea: in perseverative behaviors, past choices guide future choices. However, we did not find significant correlates of the reward-seeking variable *R*, which also guides future choices. It remains possible that MOs correlates with *R* but too weakly to detect here, for example MOs might represent the summed decision variable *P+R*, with the smaller size of *R* than *P* meaning that only *P* was individually significant. Alternatively, *R* might be stored elsewhere and only pass through MOs after the go cue, a possibility consistent with the facts that the previous trial’s interaction variable is significantly encoded after the go cue (Supplementary Figure 18), and that MOs inactivation at choice time suppresses both perseverative and reward-seeking behavior.

Our results are consistent with the hypothesis that perseveration enables rapid responses (Ashby et al., 2007; Gershman, 2020; Wood & Rünger, 2016). Reaction times were faster for repeated choices, and inactivating MOs at choice time increased reaction times. We therefore suggest that MOs activity serves to prime downstream circuits so that the moment the go cue sounds, a rapid response in the pre-planned direction is released. This priming might consist of depolarization in premotor units, or embodied changes such as altered muscle tension. When MOs is inactivated, the priming is removed, and the downstream circuit takes more time to produce an action in a direction selected close to randomly.

In summary, we identified area MOs as a region correlating specifically with a perseverative decision variable used in this dynamic reward learning task, and necessary for implementing the effect of this decision variable on choice direction and reaction time. Future work can establish which if any other areas also encode this variable especially subcortically, the mechanisms by which the decision variable is updated, and the mechanisms by which it affects downstream circuits involved in executing the actual action.

## Methods

### Initial surgery

All experimental procedures were conducted at University College London according to the UK Animals Scientific Procedures Act (1986) and under personal and project licences granted by the Home Office following appropriate ethics review.

Mice (C57BL/6, 6 male, 4 female) were implanted with a head-fixation plate over the interparietal bone and a 3d-printed guide, following the procedure described in detail in Steinmetz et al. (2019). During an initial surgery under isoflurane anesthesia (1-3% in O2), we implanted a steel headplate (size 15 x 3 x 0.5 mm; weight approximately 1g), followed by a custom guide. The guide was 3d printed from opaque polylactic acid, and when placed on an average mouse skull exposes an area from 3.5mm anterior to 5.5mm posterior to bregma, and 4.5mm left and right from bregma, allowing targeting multiple areas without a need for additional surgery, as well as protecting the probes from mouse paw movements that otherwise could break them. To implant the headplate, the surface of the skull was cleaned from the skin and periosteum and a green activator (Super-Bond C&B, Sun Medical) applied to the cleared skull, specifically to the area where the headplate was later implanted, to strengthen the connection between the bone and the cement. The skin was attached to the skull on the sides using Vetbond Tissue Adhesive (3M), and the skull surface was also covered by it, except for the area where the headplate was later implanted. The headplate and guide were then attached to the skull with dental cement (Super-Bond C&B, Sun Medical Co, Ltd, Japan).

Two layers of UV-curing optical glue (Norland Optical Adhesives #81, Norland Products) were applied to the skull and cured for 10 seconds per layer. The layers needed to be thin to prevent excessive heat. The whole procedure took around 1 hour.

After recovery, mice were treated with carprofen added to their drinking water for three days while recovering in a home cage. After this time, they were given at least an additional two days of ad lib access to water, before beginning water restriction and habituation in preparation for behavioral training.

### Task

Our task used a previously-described behavioral apparatus (Burgess et al., 2017). The mouse was head-fixed, within a half-pipe tube and covered with another half-pipe for comfort. Mouse front paws sat on a rubber wheel (LEGO, 86652 & 32019) which could be freely turned to register choices. Wheel movements were measured using an incremental rotary encoder with a resolution of 1024 increments per turn. Rewards were delivered via a silicone lickspout placed within easy reach of the mouse’s tongue. The mouse was surrounded by three video screens but no visual stimuli were displayed, just gray background.

Each trial began with a brief auditory tone (go cue, 5kHz for 100 ms), following which the mouse could turn a wheel for a liquid reward. The tone carried no information about which direction of turn would be rewarded. On each trial one of the sides was associated with 80% reward probability and 20% omission probability, and the other side was associated with 20% reward probability and 80% omission probability. The reward-probability contingencies (blocks) flipped randomly every 125-175 trials, with the duration of the first block chosen randomly between 1 and 175 trials, and the first correct side chosen at random. Block flips were not signaled to the animal. We call choices that correspond to the 80% reward probability “correct choices”, independently of whether mice are rewarded on these trials. A choice was registered when the mouse turned the wheel by 18mm in either direction. Trials did not end until this threshold was reached; there was no timeout. Upon completing a turn, the mouse received feedback (either a 1-3*μ*L drop of 10% sucrose in water or an auditory white noise) coupled with a prolonged intertrial interval. Feedback delivery was followed by an intertrial interval of random length: uniformly distributed 1 and 2 s following reward delivery, and between 2.5 and 3.5 s following reward omission. After the inter-trial interval, the mouse had to hold the wheel completely still, to within the resolution of the rotary encoder, for a “fixation period” of 200ms before the next trial began. The response time is defined as the time between the go cue and the completion of the trial.

### Behavior training

Training began with two habituation sessions (on two consecutive days), where mice were head fixed for 15 and 30 minutes, respectively. We next trained mice 1 hour per day on a deterministic reward schedule, where the correct choice was rewarded 100% of the time and the incorrect choice was never rewarded. Once mice followed block switches for at least 3 days in a row, we changed the probabilities to 90% for the correct choice and 10% for the incorrect choice. Finally, when mice were judged to perform that version well, we changed the probabilities to 80% and 20%. Mice were trained 5 days per week, and typically became proficient in the final version of the task 4-12 weeks after the start of the training. The decision on when to complete training and begin recording was subjective, based on the number of trails per session and rewards received, and all mice received at least 26 training sessions by the time recordings started (34.4 ± 4.09, mean ± S.E.M., n=10 mice). The amount of training a mouse received did not correlate with the degree of behavioral perseveration as assessed by the PR model (Supplementary Figure 4).

### Behavioural models

#### Models used

We compared the 13 models behavioral models. These were four variants of Q-learning: standard Q-learning (Sutton & Barto, 1998), optimistic Q-learning (Lefebvre et al., 2017), differential forgetting Q-learning (Ito & Doya, 2009), and the *PR* model (Miller et al., 2021); six variants of ideal observer models: standard ideal observer, ideal observer with drifts (bayes drifts), ideal observer with flips (bayes flips), ideal observer with perseveration, ideal observer with drifts and perseveration, ideal observer with flips and perseveration; and three additional models: a local matching law (Sugrue et al., 2004), a foraging agent (Constantino & Daw, 2015), and an actor-critic model (Sutton & Barto, 1998).

#### Standard Q-learning

Standard Q-learning (Sutton & Barto, 1998) is a reinforcement learning algorithm that is used to learn the optimal action-value function for an agent interacting with an environment.

The two internal variables of the Q-learning model are the action values of the left and right sides (*Q*_*L*_ and *Q*_*R*_). When a new session starts, both *Q*_*L*_ and *Q*_*R*_ are set to 0. The prediction error *δ*_*t*_ is calculated for trial *t* as a difference between the actual and the expected reward:

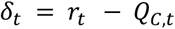

Here C represents the choice made (L or R). This calculated reward prediction error is used to update the *Q* value of the chosen action with learning rate *⍺*:

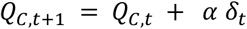

The Q value associated with the unchosen side is not updated. Decisions are made according to a softmax rule, dependent on two parameters: a bias *β*_*bias*_ and an “inverse temperature” *β*_*Q*_:

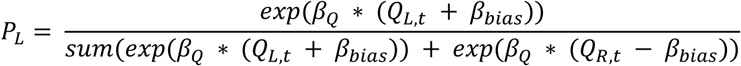

The bounds for the parameter search are: *⍺* ∈ [0,1], other parameters ∈ (−∞, ∞).

#### Optimistic Q learning

Optimistic Q-learning (Lefebvre et al., 2017) is an extension of the standard Q-learning algorithm that allows the speed of learning to differ between trials with positive and negative prediction errors:

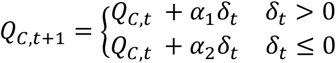

The parameter *⍺*_*l*_ the learning rate from positive prediction error and *⍺*_*2*_is the learning rate from the negative prediction error. If *⍺*_*l*_>*⍺*_*2*_, then the agent is implementing the optimistic Q-learning, modelling how people tend to incorporate good news into their world model faster than bad news (Lefebvre et al., 2017); and if *⍺*_*l*_<*⍺*_*2*_, it is pessimistic Q-learning. If *⍺*_*l*_=*⍺*_*2*_, it is standard Q-learning described in the previous section. This model has four free parameters, *⍺*_*l*_, *⍺*_*2*_, *β*_*bias*_, *β*_*Q*_. The bounds for the parameter search are: *⍺*_*l*_ ∈ [0,1], *⍺*_*2*_ ∈ [0,1], other parameters ∈ (−∞, ∞). Choices are made according to the softmax function with parameters *β*_*bias*_ and *β*_*Q*_, as above.

#### Differential Forgetting Q-learning

The Differential Forgetting Q-learning rule (Ito & Doya, 2009) allows modeling of perseveration, by causing the Q-value of the unchosen side to decay toward zero. (In standard Q-learning, the Q-value of the unchosen side does not change. It uses the following rule:

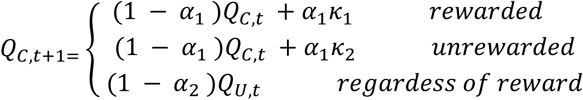

Here, *C* refers to the chosen side and U to the unchosen side (both L or R). The bounds for the parameter search are: *⍺*_*l*_ ∈ [0,1], *⍺*_*2*_ ∈ [0,1], other parameters ∈ (−∞, ∞). Choices are made according to the softmax function with parameters *β*_*bias*_ and *β*_*Q*_, as above.

#### PR model

The PR model has two decision variables: *P* which follows the law of exercise and mediates perseverative behavior, and *R* which follows the law of effect and mediates reward-seeking behavior. The perseverative decision variable *P* is initialized to zero at the beginning of every session, and updated according to:

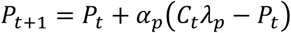

where *C*_*t*_ is the choice (left or right, coded as −1 or +1) made on trial *t*, *⍺*_*p*_ is a learning rate parameter governing the extent to which an individual choice affects *P*, and *λ*_*p*_ is a magnitude parameter governing the largest possible magnitude of *P*. This rule therefore updates *P* towards −*λ*_*p*_ following left choices and +*λ*_*p*_ following right choices, so *P* can be thought of as a running average of previous choices (Figure 1H, yellow line).

The reward-seeking decision variable R is likewise initialized to zero at the beginning of each session, and is updated according to the following rule:

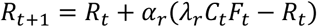

where *F*_*t*_ is the feedback (unrewarded or rewarded, coded as −1 or +1) received on trial *t*, and *⍺*_*r*_ and *λ*_*r*_ are learning rate and magnitude parameters, which can be different to the perseveration parameters *⍺*_*p*_ and *λ*_*p*_. The rule therefore updates *R* towards −*λ*_*r*_ following rewarded left choices or unrewarded right choices, and towards +*λ*_*r*_ following rewarded right choices or unrewarded left choices. Thus, *R* can be thought of as a running average of the choice-reward interaction, which provides an estimate of the correct choice (Figure 1H, orange line).

On each trial, the PR model agent chooses right over left with log odds given by the sum of the two decision variables:

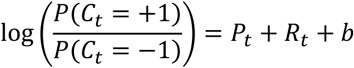

where b is a parameter governing fixed left/right bias. The overall magnitudes of P and R – determined by the parameters *λ*_*p*_and *λ*_*r*_ – therefore govern the relative influence of perseveration and reward seeking on choice. We fit the model parameters to the observed behavior of each mouse by maximum likelihood.

#### Local matching law

Hernstein’s matching law states that the rate of responding to a particular option or stimulus is proportional to the relative rate of reinforcement associated with that option or stimulus (Herrnstein, 1970). The local matching law implements this through equations are similar to the Q-learning models discussed above (Sugrue et al., 2004). The learning rate *α* reflects the speed of the action value update, the sensitivity shows the strength of the connection between the reinforcement ration and the choice ratio. Then, the Q-value update for *Q*_*L*_ can be written as follows:

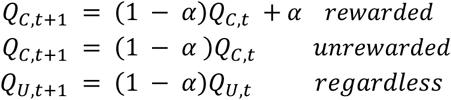

Here, *C* refers to the chosen side and U to the unchosen side (both L or R). The bounds for the parameter search are: *⍺* ∈ [0,1]. Choices are made according to the softmax function as above, but without bias terms in this case.

#### Foraging agent

The foraging agent (Constantino & Daw, 2015) explores the environment, making its decision about whether to stay with the current choice or to switch sides based on an estimate of the value of the current choice. Unlike the models above, this model only has one internal variable *V*, which estimates the value of the current choice, and resets when the agent switches sides. The value is updated according to

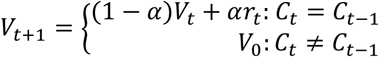

Here *C*_*t*_ = ±1 represents the choice on trial *t*. The decision is made by a logistic function with a bias for switching as well as for left/right side:

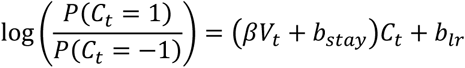

The parameters *⍺*, *β*, *b*_*stay*_, *b*_*lr*_ represent learning rate; strength of history, stay bias, and left/right bias, respectively. The bounds for the parameter search are: *⍺*_*local*_ ∈ [0,1], *V*_*global* ∈ [0, ∞), other parameters ∈ (−∞, ∞).

#### Actor-critic

Actor-critic algorithms are a family of algorithms that explicitly represent policy independent of value function (Sutton & Barto, 2018; Section 2.8). The policy structure (actor) selects actions, while the estimated value function (critic) criticizes those actions. The scalar signal of the critic drives learning of both actor and critic. In general, actor-critic algorithms allow both the actor and critic components to depend on sensory state; in the current task, with only a single state, the actor-critic model takes a particular simple form.

We considered a “forgetful” Actor-Critic model, which includes a decay rule that updates the policy variable towards zero on each trial, allowing the agent to produce adaptive behavior in an environment with changing reward contingencies (Miller et al., 2023). This model has three parameters: *⍺*_*c*_, the critic learning rate; and *⍺*_*l*_, *⍺*_*f*_, the learning and forgetting rates for the actor. It has two state variables: *V*, the critic’s expected reward; and *H*, the difference in the actor’s estimated values for actions to the left and right.

The critic learns according to the following rule:

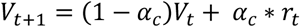

The actor updates its values according to the following rule:

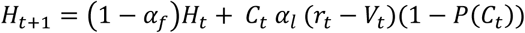

Here *C* = ±1 represents the chosen action (L or R) on trial t. 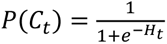 represents the probability that action *C*_*t*_ will be chosen, given by the logistic formula. Thus, the actor’s estimate of the advantage in choosing left over right is increased when it makes a left turn and receives a larger reward than the mean value *V* estimated by the critic. The term (1 − *P*(*C*_*t*_)) serves to boost the learning rate when it makes a rare choice. The search bounds for all parameters are [0,1].

#### Ideal Observer Models

An ideal observer model, also called a Bayesian filter, uses knowledge of the structure of a partially-observable dynamic environment to optimally infer the current state of that environment (Sarkka, 2013). It assumes that in each timestep the environment can be in one of several possible discrete hidden states, *h_*t*_*, and that on each trial the environment first probabilistically emits one of several possible visible observations, *o_*t*_*, and then probabilistically transitions into a new hidden state *h_*t+1*_*. An ideal observer model requires three functions containing knowledge of the environment. The first is an Initialization Function, giving the model’s belief distribution over hidden states on the first timestep: *p*(*h_*0*_*). The second is an Observation Function, giving the probability with which each observation will be produced in each hidden state *p*(*o_*t*_*|*h_*t*_*). The third is a Dynamics Function giving the probability with which the environment will transition to each possible subsequent state *p*(*h_*t+1*_*|*h_*t*_*).

The Bayesian filter model maintains a running belief distribution *B_*t*_*(*h*) over the possible current states of the environment. This distribution is initialized on the first trial to match the Initialization Function, and then updated after each trial in two steps. The first step makes use of the Observation Function and computes a posterior probability over *h_*t*_* conditional on the observation *o_*t*_* using Bayes’ theorem. The second step makes use of the Dynamics Function to compute a belief over *h_*t+1*_* marginalizing over this posterior.

### Optimal Agent

We first consider the optimal agent, built around an ideal observer model with full knowledge of the true rules of the task. Specifically, it knows that the environment can be in one of 350 possible states: The current high-reward choice can be either left or right, and there can be between 1 and 175 trials remaining in the current block. We can therefore represent the agent’s belief distribution as a pair of vectors of length 175, the sum of whose elements is together 1. The Initialization Function is uniform, as at the beginning of the session it is equally likely that the rig will be in any of these states. We therefore initialize the vectors uniformly at the start of each session by setting each element to 1/350.

The Observation Function is given by the reward contingencies: if the mouse selects the high-reward choice, reward is delivered with probability 80%, and if it selects the low-reward choice, reward is delivered with probability 20%. We therefore update each element of the belief distribution after each trial using Bayes’ theorem and renormalize.

The Dynamics Function is given by the tasks’ block switching rules: on each trial the “trials remaining in block” variable is decremented by one, until it reaches 0 in which case the “high-reward” variable is flipped and “trials remaining in block” is sampled uniformly between 125 and 175. We therefore move probability mass down the vectors by one slot, such that the belief on trial *t* that there are exactly *n* trials remaining in the block is equal to the belief on trial *t*+1 that there are exactly *n*-1 trials remaining. The belief from the final slot (belief that this was the final trial in the block) is redistributed uniformly over the first 50 slots in the alternate vector (e.g. if the model had belief 50% that this was the final trial in a left-better block, this will add 1% belief to each element between “125 trials remaining in a right-better block” and “175 trials remaining in a right-better block”).

In this way, the ideal observer optimally updates a belief distribution over the true hidden states of the experiment. It can use this distribution to select actions, marginalizing over the number of remaining trials to compute an overall belief that it is currently in a left-better vs a right-better block. We convert this ideal observer into an optimal agent by having it deterministically select the action which it believes is currently high-reward.

### Noisy Ideal Observer

We define the noisy ideal observer agent as containing the same ideal observer model as the optimal agent, but selecting actions using a softmax decision rule with fit parameter β_*b*_:

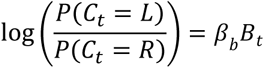

Where B_*t*_∈(0, 1) is the ideal observer’s belief on trial t that the left action is currently the highly rewarding action. The noisy ideal observer agent is therefore characterized by the one free parameter β_*b.*_

### Bayesian filter assuming probabilistic reversals (Bayesian flips)

The Ideal Observer agents make use of detailed knowledge of the dynamics of our particular task that it is unlikely a mouse would be able to perfectly infer. We therefore considered a Bayesian filter agent assuming a more general task structure: that there are two discrete blocks, each with a high-reward and low-reward side delivering reward with distinct probabilities, and that these blocks change unpredictably over time.

This set of assumptions corresponds to a belief in a single hidden variable: the identity of the current high-reward side. The belief state of the agent on each trial is therefore characterized by a single number B_*t*_ giving its current belief that the left action is high-reward. We initialize this belief to 0.5 at the beginning of each session, and update it following each trial. The Observation Function involves two free parameters P_*high*_ and P_*low*_, giving the model’s assumption about the reward probabilities on the high-reward and low-reward side. The Dynamics function involves one free parameter P_*flip*_ giving the models assumption about the probability with which the block will flip on each trial.

The Bayesian filter assuming probabilistic reversals therefore has four free parameters: β_*b*_, P_*high*_, P_*low*_, and P_*flip*_.

### Bayesian filter assuming time-varying probability changes (Bayesian drifts)

The Bayesian filter assuming probabilistic reversals still contains structural knowledge about our task that might not be available to the mouse: that reward probabilities change discretely and reverse between a particular set of fixed values. We therefore also considered a Bayesian filter agent assuming an even more general task structure: that each action is associated with a separate reward probability, and that the value of each of these probabilities on each trial tends to resemble its value on recent past trials.

This set of assumptions corresponds a belief distributions over the full set of probabilities between 0 and 1 for each action. We use *θ*_*left*_ and *θ*_*right*_ to parameterize the reward probability on the left and on the right port, and B_*right,* *t*_ and B_*left,* *t*_ to indicate the observer’s belief at time t over these probabilities. We approximate these distributions numerically as histograms, allowing *θ* to take 200 discrete values equally spaced between zero and one. This means that the first element of B_*left,* *t*_, for example, corresponds to the observer’s current belief that the reward probability of the left action is currently between 0 and 0.5%

Belief distributions are initialized to be uniform at the start of each session. After each trial, we first update the vector corresponding to the chosen action using the Observation Model given by the average probability in each bin (for example the first element is updated on the assumption that rewards are given in this state 0.25% of the time). We next update both vectors on the assumption that probabilities can change over time, by convolving them with a gaussian kernel whose standard deviation (σ) is a free parameter corresponding to the observer’s belief about the plausible rate of environmental change. Probability mass which would be spread by this convolution to values less than zero is placed instead into the lowest bin (0% to 0.5%), and likewise mass which would be spread to values greater than one is placed into the highest bin (99.5% to 100%).

To make choices, the observer marginalizes these distributions, converting a belief distribution over all possible values of the generative reward probability into a single number indicating the probability with which the observer expects a reward if it selects this action:

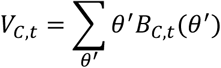

Where the index C can take the values L or R. This agent selects actions according to the difference in reward probabilities between the two actions:

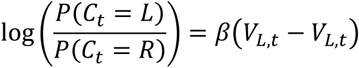

This Bayesian filter agent therefore has two free parameters: σ and β

### Ideal Observers with Perseveration

For each of the three Bayesian filter agents (Noisy Ideal Observer, Bayesian Flips, Bayesian Drifts), we also define and fit “with perseveration” versions, which are mixture models comprised of the Bayesian filter alongside a perseverative agent as defined in Equation (1):

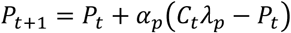

The decision rule of these agents is a mixture incorporating both the Bayesian filter’s decision variable as well as a perseverative decision variable:

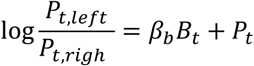

These “with perseveration” versions therefore have all of the free parameters of the associated Bayesian filter model, as well as the two additional free parameters α_*p*_ and λ_*p*_.

#### Fitting procedure

The parameters of all behavioral models were fit by maximum likelihood. Specifically we used MATLAB’s fmincon function, with parameters: maxfunevals = 4000, maxiter = 2000, algorithm = ‘interior-point’. For each parameter of each model, we manually specified a feasible range for the parameter search, and selected a starting value randomly in this interval. We repeated the fmincon procedure until it completed successfully three times, and recorded the parameter fits from the run that has the best likelihood of these three runs. The runs were consistent: for all but three models the difference in log-likelihood between runs was no larger than 1e-5. Among the remaining three models, two were non-perseverative (Bayesian flips and a foraging agent). For these models, we ran the fitting procedure 20 times and the log-likelihood for every run was lower than the log-likelihood of any perseverative model. For the remaining model (Bayesian flips with perseveration) we ran the fitting procedure 20 times; it had high-varying log-likelihood, but in all cases it was higher than the log-likelihood of all non-perseverative models.

For model comparison, we use a cross-validation approach that allowed for differences in strategy between mice. We fit each model’s parameters on odd sessions of each mouse separately, and evaluated the model by log likelihood on the even sessions using parameters fit to the same mouse; the model parameters were thus assumed to differ between mice, but not between sessions of one mouse. The quality of fit for each mouse was obtained by dividing the total log likelihood by the total number of trials, then exponentiating to yield a geometric mean likelihood.

#### Trial-history regression

To gain a better understanding of the effect that MOs suppression has on choices, we compared trial history regression analyses (Lau & Glimcher, 2005; Lee et al., 2004) on trials with and without optogenetic suppression of MOs activity. The regression included separate terms for the choice, feedback and their interaction for past trials trying to predict the choice on the current trial. There are thus separate weights for each of the two possible choices (left, right), each of the two reward outcomes (rewarded, unrewarded), and each of the possible interactions (left rewarded or right unrewarded vs. right rewarded or left unrewarded), for each time lag considered. The pattern of trial-history regression weights provide information on the decision strategy used by the mice, without reference to explicit psychological models (Miller et al., 2022).

The full equation for trial history regression is:

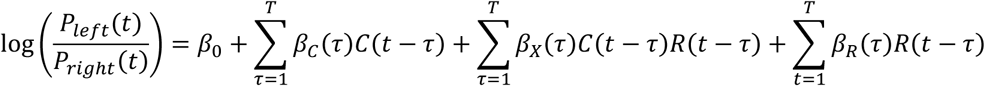

This equation expresses the log odds choice on trial *t* as a function of past choices and rewards. *C* indicates past choices, taking the value 1 for left choice trials (either rewarded or not) and −1 for right choice trials. Positive values of β_*c*_, therefore, indicate a tendency to repeat past choices. *R* indicates past rewards, taking the value 1 for rewarded choices (either left or right) and −1 for non-rewarded choices. Positive values of β_*x*_, therefore, indicates a tendency to repeat choices that have led to rewards and to switch away from choices that have not. Positive values of β_*r*_ would indicate a tendency to select the left action following rewards, regardless of the choice that preceded those rewards. T is a hyperparameter that governs the number of past trials that go into the model to predict the upcoming choice, set to 10 for the current analyses. The weights *β* were fit by maximum likelihood using MATLAB’s glmfit function with default options.

### Secondary surgery for electrophysiological experiments

After mice were fully trained in the task, they were retrained in the recording rig, which typically required additional 3-10 days.

On the first day of recordings, mice were anesthetized (1-3% isoflurane in O2) and a <0.5mm craniotomy was made over the desired location with a dental drill (coordinates: 1.8AP, ±2ML for targeting DMS; −1.6AP, ±0.9ML for targeting the hippocampus; 2.2AP, ±0.9ML for targeting MOs). The location of the craniotomies and the desired probe angles were determined using custom software (https://github.com/petersaj/AP_scripts_cortexlab/tree/master/allen_interface). A thin layer of Kwik-Cast (World Precision Instruments) was applied on top of the craniotomy to protect it. The animals were allowed to recover for at least 3 hours before the recording began. For each additional recording location, the procedure was repeated, with a maximum of three locations per mouse.

### Electrophysiological recordings

Neuropixels probes, either 1.0 version 3B or 2.0, were held by metal rods and gradually lowered into the brain using a motorized micromanipulator (uMP-4, Sensapex Inc.), with an approximate speed of 10 micrometers per second, piercing the dura mater. To allow later track localization, prior to insertion probes were coated with a solution of DiI (ThermoFisher Vybrant V22888, V22886 or V22885) by holding 2μL in a droplet on the end of a micropipette and touching the droplet to the probe shank, letting it dry, and repeating until the droplet was gone, after which the probe appeared colored. After smooth descent to the final depth, probes were left untouched for at least 30 mins before recording. The plastic well was filled with physiological saline for the duration of the recording, prevent drying. Recordings were performed using a PXI system (NI PXIe-1071) and spikeGLX software. The recorded data were spike sorted using Kilosort 2.5 (Pachitariu et al., 2023) and then manually curated to remove noise clusters with phy (https://github.com/kwikteam/phy). Units were considered “noise” and removed from further analysis if they had non-physiological waveform shape or pattern of activity across channels. No further manual processing was performed. We use the term “neural activity” throughout the manuscript to mean spike counts in a window of fixed length.

### Histology

Following recordings, mice were transcardially perfused following standard procedures. Briefly, they were administered a terminal dose of pentobarbital intraperitoneally, followed by a toe-pinch to ensure anesthesia. The thoracic cavity was opened, the right atrium was incised, and a solution of phosphate buffered saline (PBS) followed by 4% formaldehyde (Thermofisher 28908) in 0.1M PB pH 7.4 was perfused through the left ventricle. The brain was carefully dissected out and post-fixed in the same fixative for a minimum of 24 hours at room temperature. After washing, the tissues were stored in PBS at 4°C for no longer than three weeks before being imaged *in toto* with serial section two-photon microscopy (Economo et al., 2016; Ragan et al., 2012). The probe and cannula locations were then estimated using MATLAB software (https://github.com/petersaj/AP_histology).

### Unit alignment

The three-dimensional location of each recorded unit was aligned to the Allen common coordinate framework atlas (Q. Wang et al., 2020) using custom MATLAB code (http://github.com/petersaj/AP_histology). For the purposes of analysis, we labelled units as belonging to “MOs” if they were in Allen atlas region “Secondary motor area”; to “mPFC” if they were in “Prelimbic area”, “Infralimbic area” or “Anterior cingulate area”; we did not aim for higher specificity to avoid possible mislabeling due to probe depth. All units in Allen region “Caudoputamen” were assigned a label “DMS” due to their dorsomedial location. We assign a label “Dorsal hippocampus” to any of the units in CA1, CA3 or dentate gyrus. We assign the label “Orbital areas” to any of the units in the orbital area (including lateral and medial subdivisions). We assign a label “Olfactory areas” to any of the units in the olfactory area (including main olfactory bulb, piriform area and others).

We recorded activity from more brain areas than analyzed in the paper, however the statistical methods we used require at least 6 sessions with at least 30 neurons each, therefore we did not apply these analyses to these all recorded areas (all areas recorded and number of sessions per region listed in Supplementary Figure 6).

### Neural data analysis

#### Predicting a behavioural variable from neural activity

To assess whether a behavioral variable is coded by the neural activity, we used the session permutation method (Harris, 2021). This analysis asks whether neural data better predicts behavioral variables recorded simultaneously on the same session, than behavioral variables recorded on another session. This tests the null hypothesis that the full history of neural activity of a session is independent of the full sequence of behavior of that session, but makes no assumption of independence of neural or behavioral variables between trials, or of independence between cells.

The session permutation uses a test statistic ∑_*s*_ *ρ*(***A***_*s*_, _*s*_), where ***A***_*s*_ and _*s*_ represent the histories of neural activity and behavior on session *s* encoded into arrays with *N*_*trials*_rows, and *ρ* is a scalar measure of correlation to be defined below. It compares this test statistic to a null ensemble {*ρ*(***A***_*s*_, _*rr(s)*_): *π* ∈ Π}, where Π is a set of 1000 random permutations of the session labels. To use the method, all sessions must be truncated to the same length. Here we use all sessions containing at least 30 neurons in a specified area, and truncate each on to have the same number of trials as the shortest of these sessions.

The correlation measure *ρ* was different for different types of behavioral data considered. When correlating neural activity with a continuous behavioral variable (such as the decision variable of the PR model), we used sparse linear regression. For every behavioral session, we found the 20 neurons with largest squared correlation with the behavioral variable, predicted the behavioral variable on each trial from these neurons’ activity using unpenalized linear regression, assessed performance for each session with mean squared error (MSE), and summed across sessions to get the test statistic We then generated 1000 random permutations of the sessions, and recomputed the statistic 1000 times, but now predicting the behaviour on session *s* using the neural activity of session *π*(*s*). We deemed the correlation significant if the test statistic is below the first percentile of the null ensemble.

To predict a binary variable (such as left/right choice), we applied the same procedure but using logistic regression, with fit quality evaluated by log likelihood instead of MSE. We deemed the result is significant if the real likelihood was above the 99^th^ percentile of the likelihoods of shuffles.

To verify that any differences did not arise from different numbers of neurons recorded in different areas, we ran an analysis on equally subsampled set of neurons. We selected all sessions containing at least 60 neurons in a specified area and as before kept the first *L* trials of every session. Then, we randomly subsampled either 10, 20, 30, 40, 50 or 60 neurons and repeated the analysis above. We deemed MOs to be significantly different from other areas for a specific neuron count if the mean difference between the real and shuffled sessions in MOs was above the 99^th^ percentile of such differences in 5 other regions.

#### Accounting for other behavioural variables

Since the decision variable is correlated with choice, we tested for a partial correlation of neural activity with the decision variable given the choice: does neural activity predict the decision variable even if a common correlation with choice is accounted for?

To do so without assuming independence of trials or neurons within a session, we used an adaptation of the session permutation method that tests for partial correlation (Harris & Yuan, 2024). Again this requires all sessions to have the same length, so we took only sessions containing at least 30 neurons in the brain area of interest, and truncated each to have the same number of trials *L* (the length of the shortest session).

To perform partial correlation, we first construct a *L* × *L* matrix Π that projects out the choice variable from each session. To do so, we construct a matrix *D* of size *L* × *N*_*sessions*_ containing the sequence of choices for each session. We then apply singular value decomposition to obtain a matrix *U* of the same size as *D*, with orthogonal columns spanning the same subspace as the columns of *D*, and define Π = *I* − *UU*^T^. Denoting the L-dimensional timeseries of behavioral values for session *s* as *b*_*s*_, we next predict Π*b*_*s*_ from the neurons recorded in session *s* using a sparse method: we find the 20 neurons that have the highest correlation with Π*b*_*s*_ and fit an unpenalized linear regression from their fixation period activity to predict Π*b*_*s*_. We compute the test statistic as the average of the MSE of this regression over all sessions. To gauge significance, we compare this test statistic to a null ensemble in which the behavioral variables, but not the neural activity, has been permuted across sessions. Because this session permutation does not affect the projection matrix Π, it yields a test for partial correlation of the behavioral variable with neural activity, allowing for a common effect of choice.

#### Correlation of reaction time and neural activity in MOs

For every session containing at least 30 neurons in MOs, we compute the firing rate of each neuron in the 200ms fixation period prior to the go cue, for every trial. We select the 20 neurons whose fixation-period activity best correlates with the left/right choices (assessed by Pearson correlation). We then fit an unpenalized logistic regression model to predict the left/right choices on each trial from the resulting 20-dimensional vectors. We then compare summed neural activity arising from this model (prior to passing through the logistic function) on each trial to the animals’ reaction time on that trial.

#### Causal analysis of effect of reward or interaction on neural activity

We took advantage of the fact that rewards were delivered at random to assess whether reward and reward-choice interactions had a causal effect on neural activity in the subsequent fixation period. To do this, we computed a test statistic measuring the correlation of the reward (or interaction) on each trial with fixation-period activity of the next trial and averaged it across sessions; then compared this statistic to a null ensemble obtained by randomly regenerating the reward sequence using the same rules as the original session.

The test statistic was defined by sparse logistic regression. For each session, we found the 20 neurons whose activity had the highest correlation with the previous trial’s reward (or interaction), and applied unpenalized logistic regression to predict the reward (or interaction) from their activity. We defined the test statistic as the average log likelihood of this prediction over trials. We compared this test statistic to a null ensemble obtained by repeating 1000 times with regenerated reward sequences, choosing the 20 best neurons and prediction weights again each time. We infer that the correlation is significant if the test statistic is above the 99^th^ percentile of the null distribution. The analysis was run multiple times, computing activity for different time bins.

### Optogenetics

To assess whether the neural activity in the MOs and mPFC plays a causal role in the task, we applied transient optogenetics inactivations to these areas. Ai32 x PV-Cre (Ai32 [Jax #012569, RRID:IMSR_JAX: 012569] x PV-Cre [Jax #008069, RRID:IMSR_JAX: 008069], 4 male, 9 female) mice were implanted with glass cannulas (Doric MFC_200/230-0.48_2mm_MF1.25_FLT) bilaterally, using coordinates 2.2AP, 0.9 ML (figure 6, Supplementary Figure 21). In these mice, blue light excites PV interneurons, thereby inhibiting the pyramidal neurons (Cardin et al., 2010; Rikhye et al., 2021). To address a possible concern about the inactivation spreading beyond the target brain region (Li et al., 2019), the dataset consists of 6 mice in which the cannulas were installed on the surface of the brain targeting MOs (MOs group, 2 male, 4 female, figure 6 A) and 7 mice in which the cannulas were implanted in the same AP and ML location, but 1mm deeper than the first group, and slightly angled (mPFC group, 2 male, 5 female, figure 6 A). This is because the goal of the experiment with the MOs group was to see if inhibiting MOs specifically affects choice behaviour. The mPFC group thus also served as a negative control for the MOs group: an effect that is specific to the MOs group could not arise due to deeper areas being inactivated. Due to a health issue, one mouse of the mPFC group was only used in the “half session” experiment but not in the “random inactivation” experiment.

We ran two sets of experiments (figure 6 B, C, figure 7A, B) with the same population of mice. In the first experiment, light was delivered on 20% of randomly chosen trials, starting with the go cue until the animal moved a wheel or for 4.8s, whichever was faster. On a further 20% of randomly chosen trials, light was delivered starting at the time of feedback for 1.6s. Since the two stimulations are independent of one another, inhibition happens during both the choice period and the outcome period on 4% of trials. Light consisted of a train of 10ms blue (473 nm) laser pulses separated by 30ms gaps, delivered through an optic fiber (NA = 0.22, Doric Lenses) using a diode laser (Omicron LuxX 473-100). The laser power was 63-110 mW/*mm*^2^, measured at the tip of the patch cable that was then attached to the implanted cannula before the start of every experiment.

In the second set of experiments, stimulation was not applied on random trials. Instead, we randomly chose either the first part of the session (from trial 1 until trial 250), or the second half (from trial 251 until the end of the session) and applied the stimulation on every trial of the chosen session part at the time of feedback delivery for 1.6s.

### Optogenetic data analysis

#### Analysis of the half-session opto experiment

We constructed learning curves by computing the proportion of choices equal to the correct choice, as a function of trial number relative to the block switch, averaging over switches in all sessions. We did so separately for block switches during the laser stimulation half-session and in the other half-session. The test statistic was defined as the difference between these two curves. To assess whether the difference between the two conditions is significant, we constructed a null ensemble of 100 randomizations by recalculating this statistic after resampled the half of each session where the laser was on, according to the same rule as for the real experiment.

### Reaction time and firing rate relative to block switch

To estimate mean reaction time for different trials relative to block switch (Supplementary Figure 5A), we first identified all choice switch trials (i.e. trials N where *C*_*N*_ ≠ *C*_*N-l*_). To estimate the mean reaction time n trials before the switch, we then identified the subset of these switches for which all choices from trial N-n to trial N-1 where identical; this excluded cases where choice switched twice between trial N-n and trial N. Similarly, to estimate mean reaction times n trials after the switch, we identified the subset of switches for which all choices from trial N to trial N+n were identical. For each session and each n, we then took the median reaction time of these subsets of trials (N+n or N-n for each n), and subtracted the overall median reaction time for the session. We identified statistical significance by a t-test that this difference had mean 0.

To investigate the relationship of reaction time with the perseveration or reward-seeking decision variables of the PR model (Figure 2D), we computed a mean of the binned reaction times at 8 points equally distributed from –*λ*_*p*_ to *λ*_*p*_ and subtract the mean of these. We deem the difference significant when it’s different from 0 according to Wilcoxon signed rank test with 5% significance level.

## Acknowledgements

We thank Charu Reddy and Matteo Carandini for experimental help and discussions. This work was supported by Wellcome Trust grants 223144, 205093 and ERC grant 694401 to K.D.H.

## Author contributions

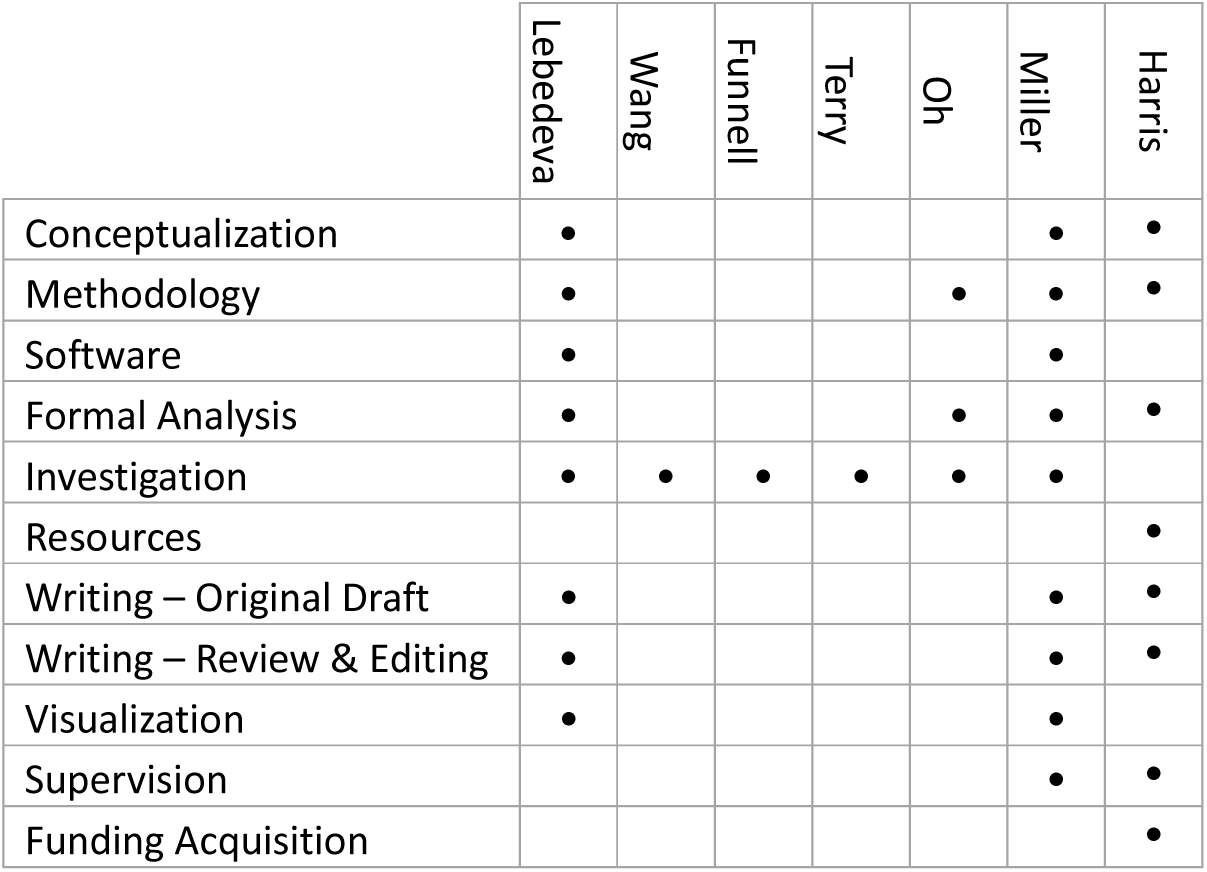

## Competing Interests

The authors declare no competing interests.

## Supplementary figures

**Supplementary figure 1.**
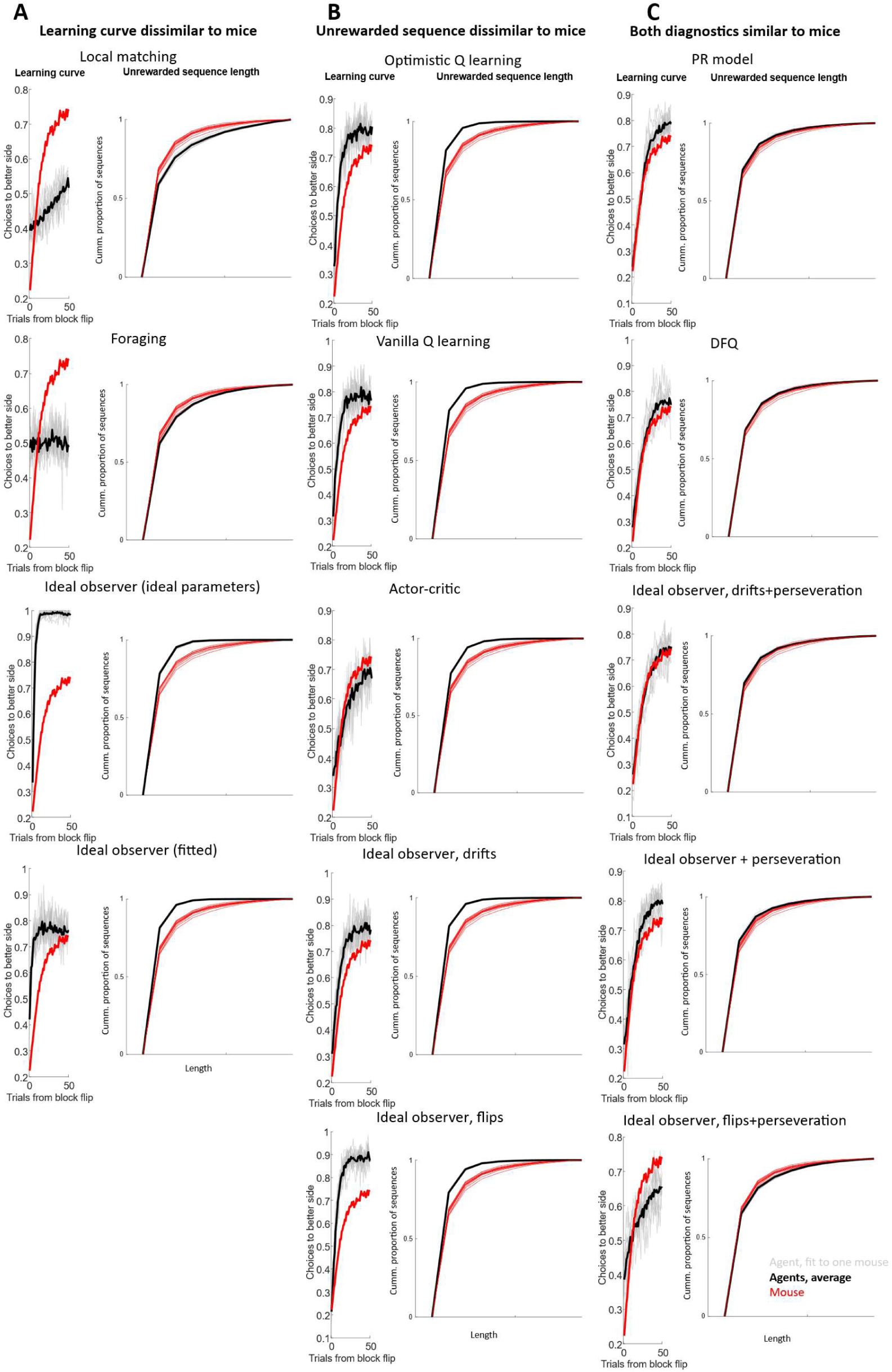
Diagnostics for all models. Two behavioural diagnostic plots are shown for each model: the learning curve (fraction of choices to the correct side as a function of time since a block switch, as in Fig. 1D); cumulative proportion of the lengths of unrewarded sequences, as in Fig. 1E. Grey lines, data for individual fits; black line, average. A, Models in which the first diagnostic is dissimilar to mice. B, Models in which the first diagnostic is dissimilar to mice. C, Models in which both diagnostics are similar to mouse diagnostics. Note, all models in C are perseverative.

**Supplementary figure 2.**
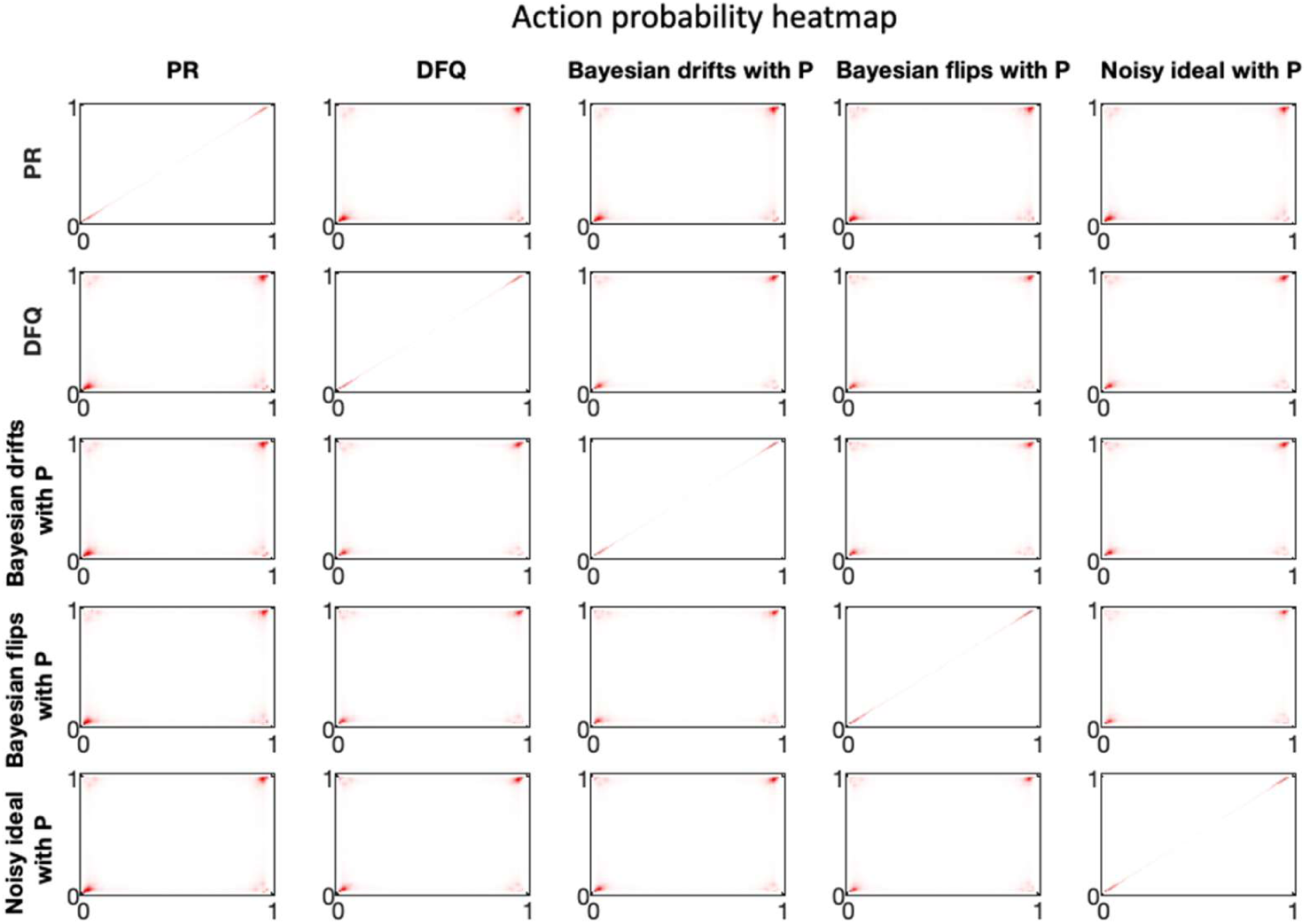
Similarity of choice predictions for the 5 winning models. Each panel shows a pseudocolor histogram summarizing the frequency that every pair of choice probabilities was predicted by two models, across all trials, sessions, and mice. For example, the intensity in the top left corner of the top right plot shows the frequency of trials on which the PR model predicted p(Right)=0 and the ideal observer model predicted p(Right)=1.

**Supplementary figure 3.**
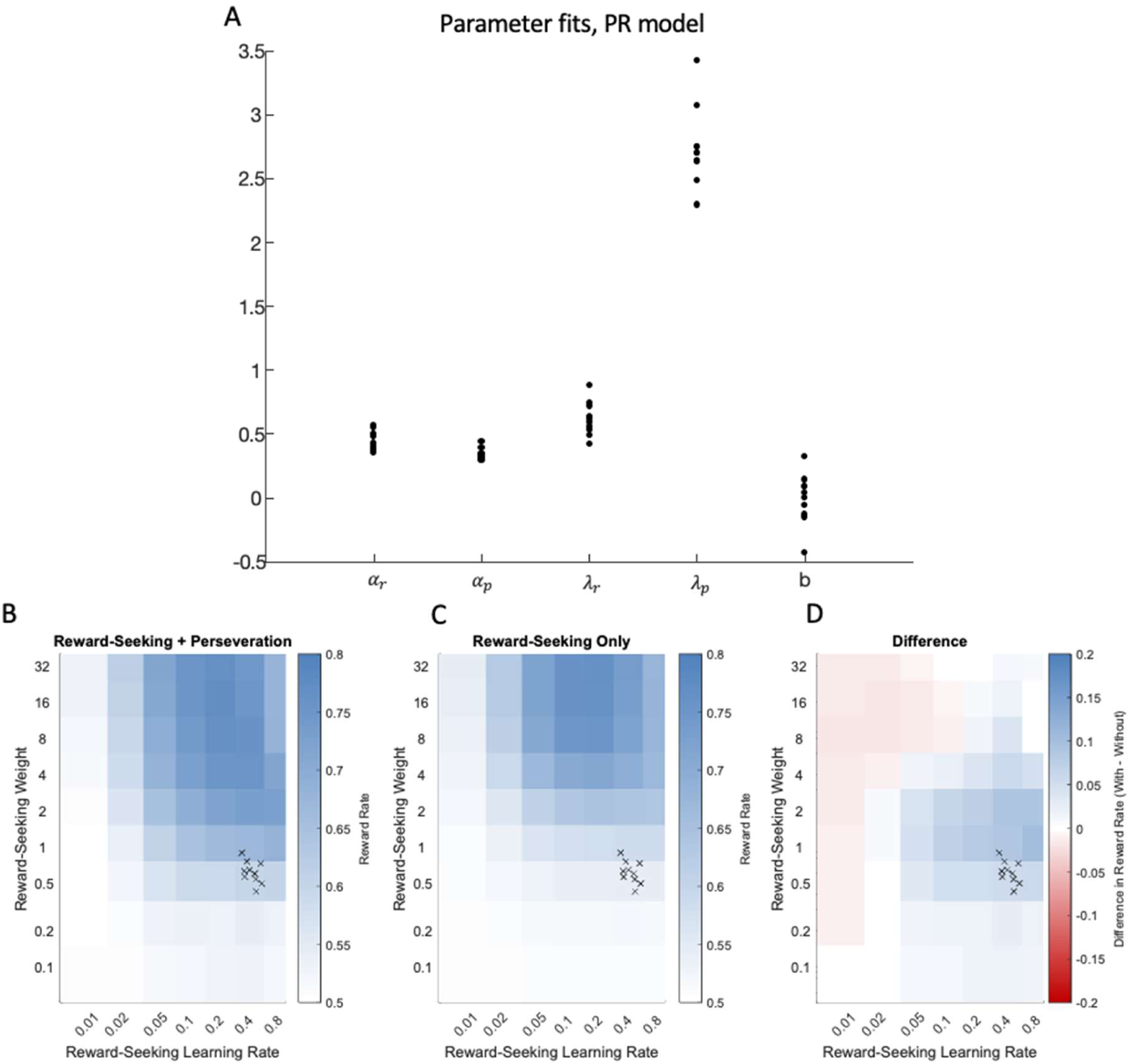
Fit parameters: PR model. A, The parameter fits for the PR model for each mouse. B, reward rate obtained in simulations of the PR model, varying the reward seeking weight *λ*_*r*_ and reward learning rate *⍺*_*r*_ parameters, while maintaining the mean values of the perseveration variables *⍺*_*p*_ and *λ*_*p*_ fit to mouse behavior. Crosses indicate the values of *⍺*_*r*_ and *λ*_*r*_ fit to mouse data. The highest reward rates are obtained at much larger values of *λ*_*r*_ and slightly lower values of *⍺*_*r*_ than those used by the mice. C, same but for simulations with no perseveration (*⍺*_*p*_ = 0), indicating the reward-rate that can be obtained by a reward-seeking agent on its own. D, difference between the left and middle panels. Including a perseverative agent with mouse-like parameters helps, rather than hinders, reward rate over much of the range.

**Supplementary figure 4.**
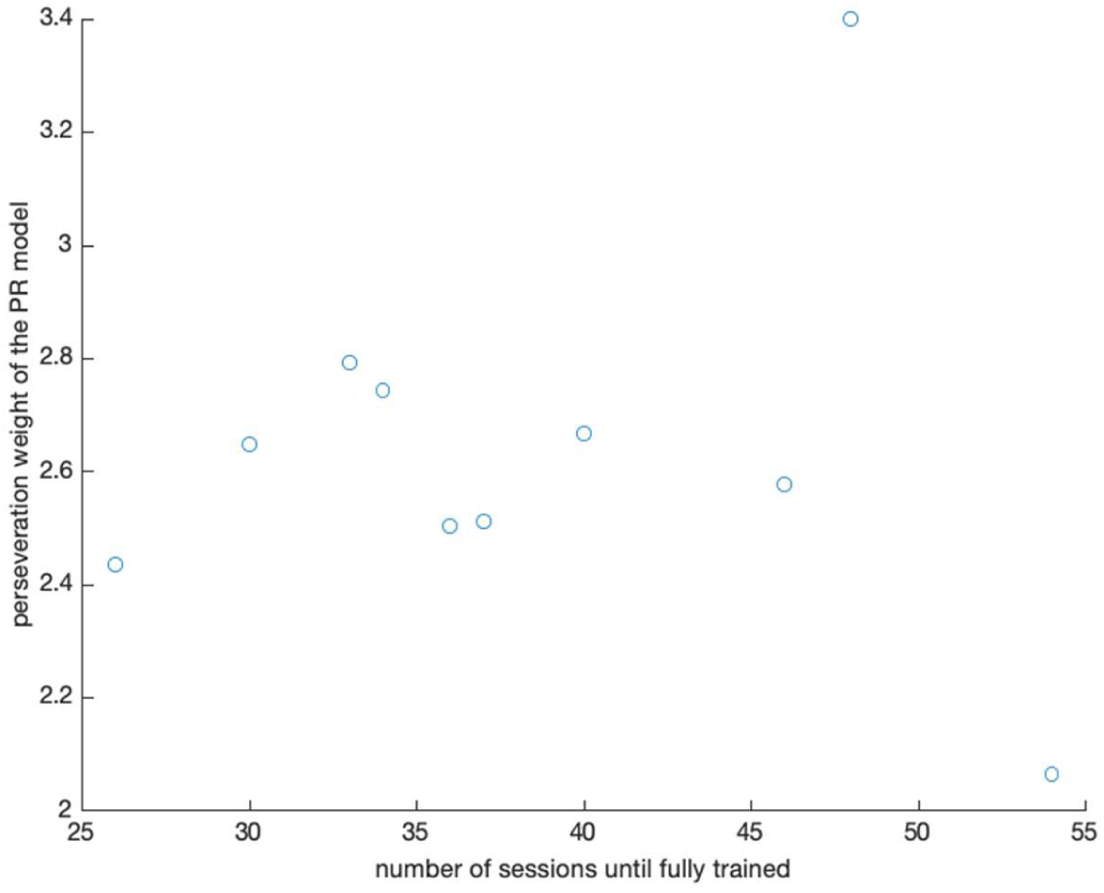
Number of sessions until animal was fully trained on the task against the perseveration weight in the PR model based on this animal’s performance. The decision to call an animal fully trained was made by the experimenter subjectively and was based on the number of trials in each session, as well as the reward received.

**Supplementary figure 5.**
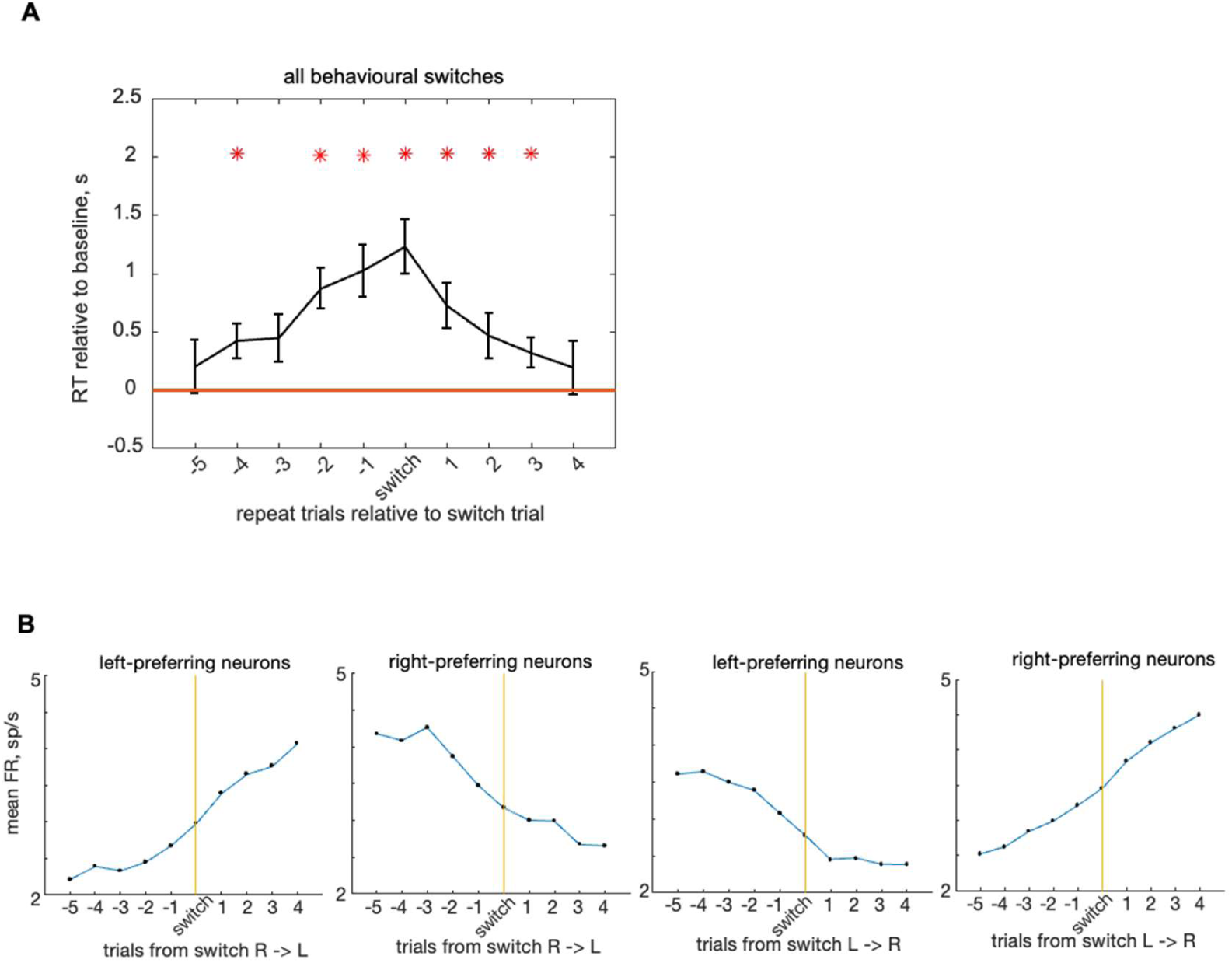
Changes in response time and neural activity in MOs relative to behavior switches. **A**, median response time as a function of the number of trials relative to the choice switch, minus overall median response time. An x-axis value of n indicates median reaction times for all trials exactly n trials following a switch, without a second intervening switch. Thus the trial following the sequence LRRR would go into bin 3 only, while the last trial of sequence LRLR go into bin 1 only. Red stars indicate a significant difference from 0. **B**, Mean fixation-period activity of the 20 units that correlate best with either right or left choice, as a function of the number of trials relative to a choice switches from right to left or from left to right, on average across all MOs recordings.

**Supplementary figure 6.**
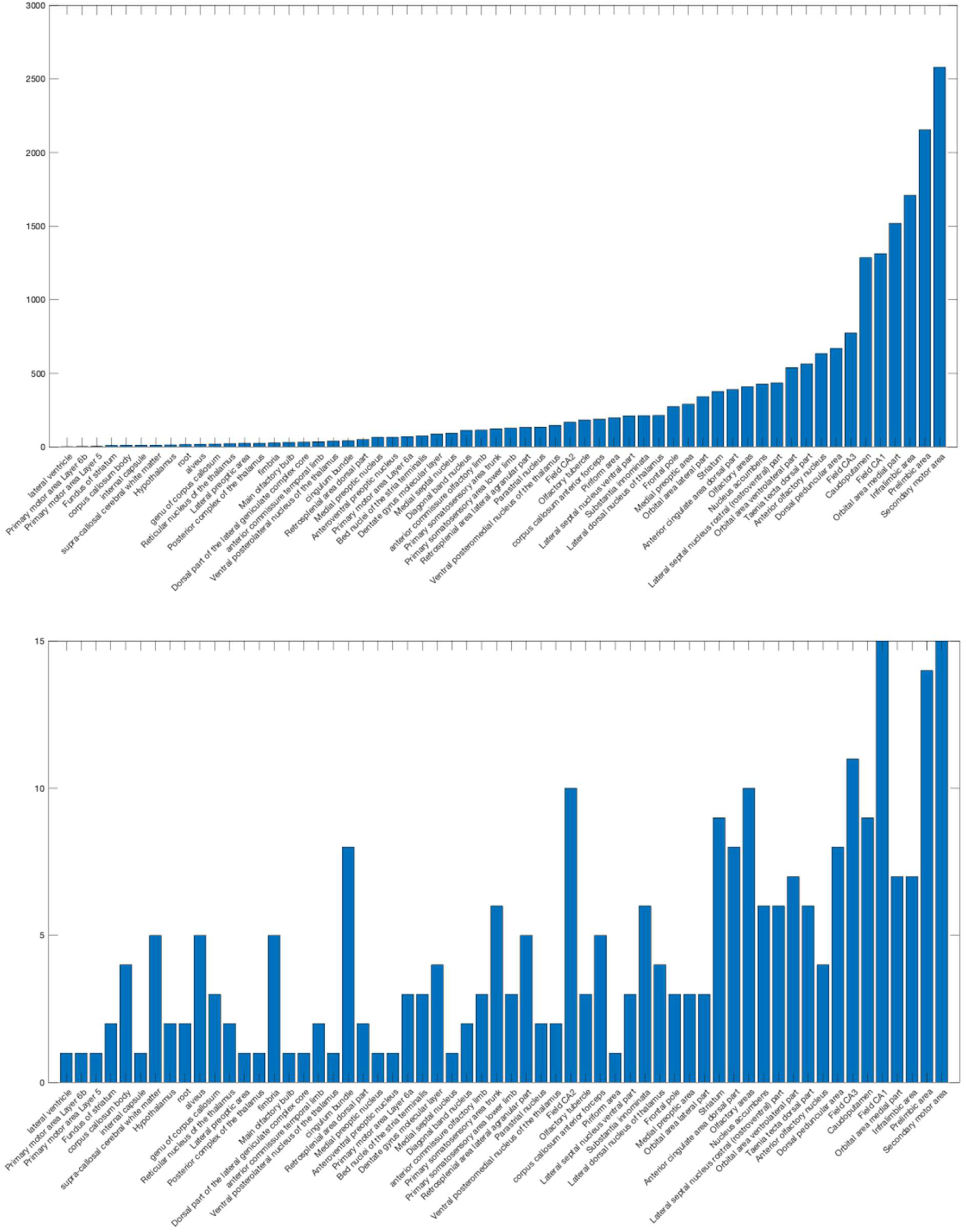
Top, number of units recorded in each area. Bottom, number of recording sessions per area.

**Supplementary figure 7.**
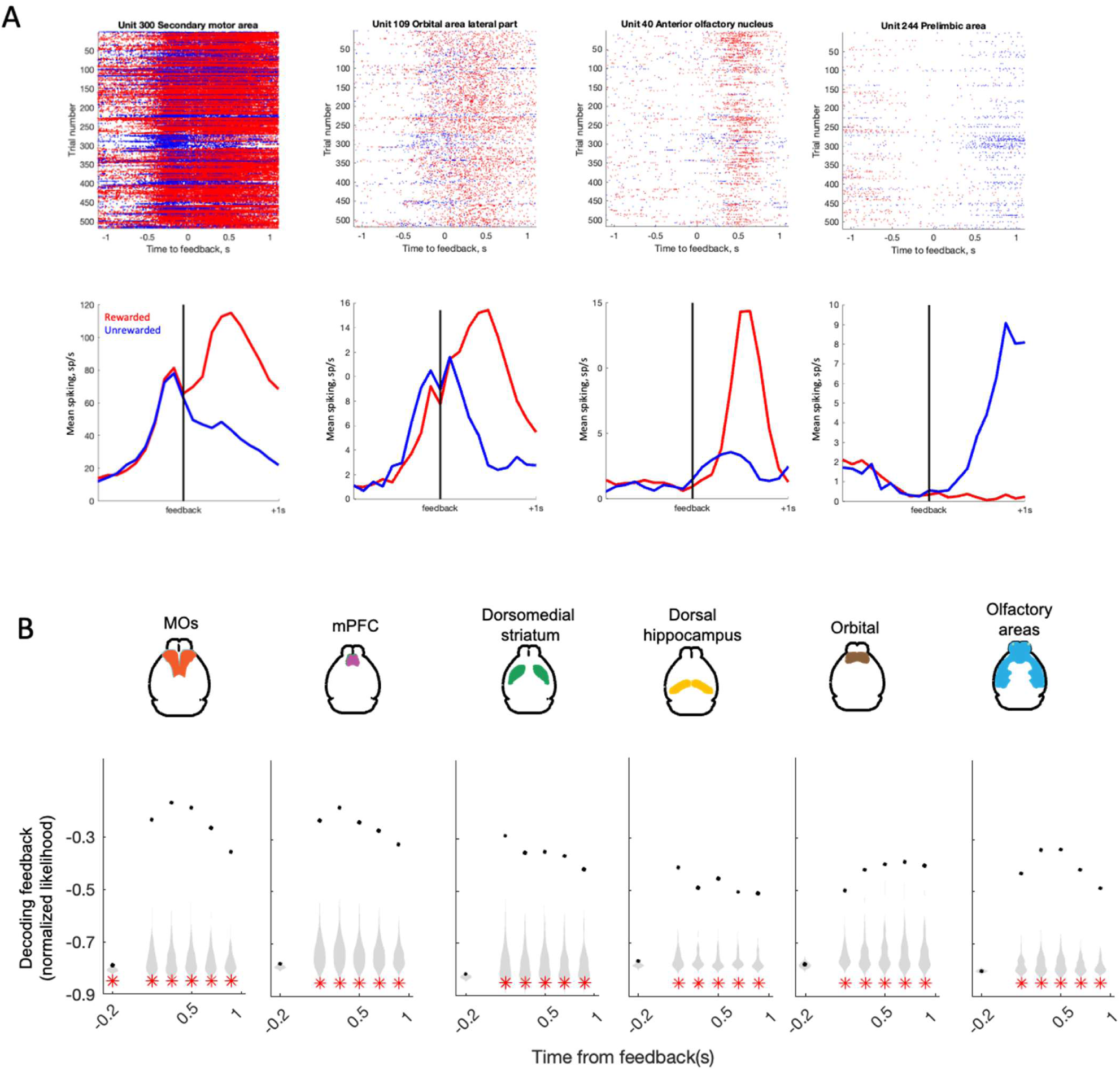
**A,** Activity of example units aligned to the feedback time, separated into rewarded (red) and no rewarded (blue) trials. Top: raster plots for four example units, trials ordered chronologically and colored by trial type. Bottom: peristimulus time histograms for the same units. Activity of each trial is aligned to the feedback time on that trial. **B,** Decoding of feedback from population activity. X-axis: time epoch; 0-200 ms before feedback, and subsequent 200 ms epochs from feedback onset. Black dots: mean prediction of actual value across sessions; gray violins, null distribution from session permutation. Red stars: p<0.01, session permutation test.

**Supplementary figure 8.**
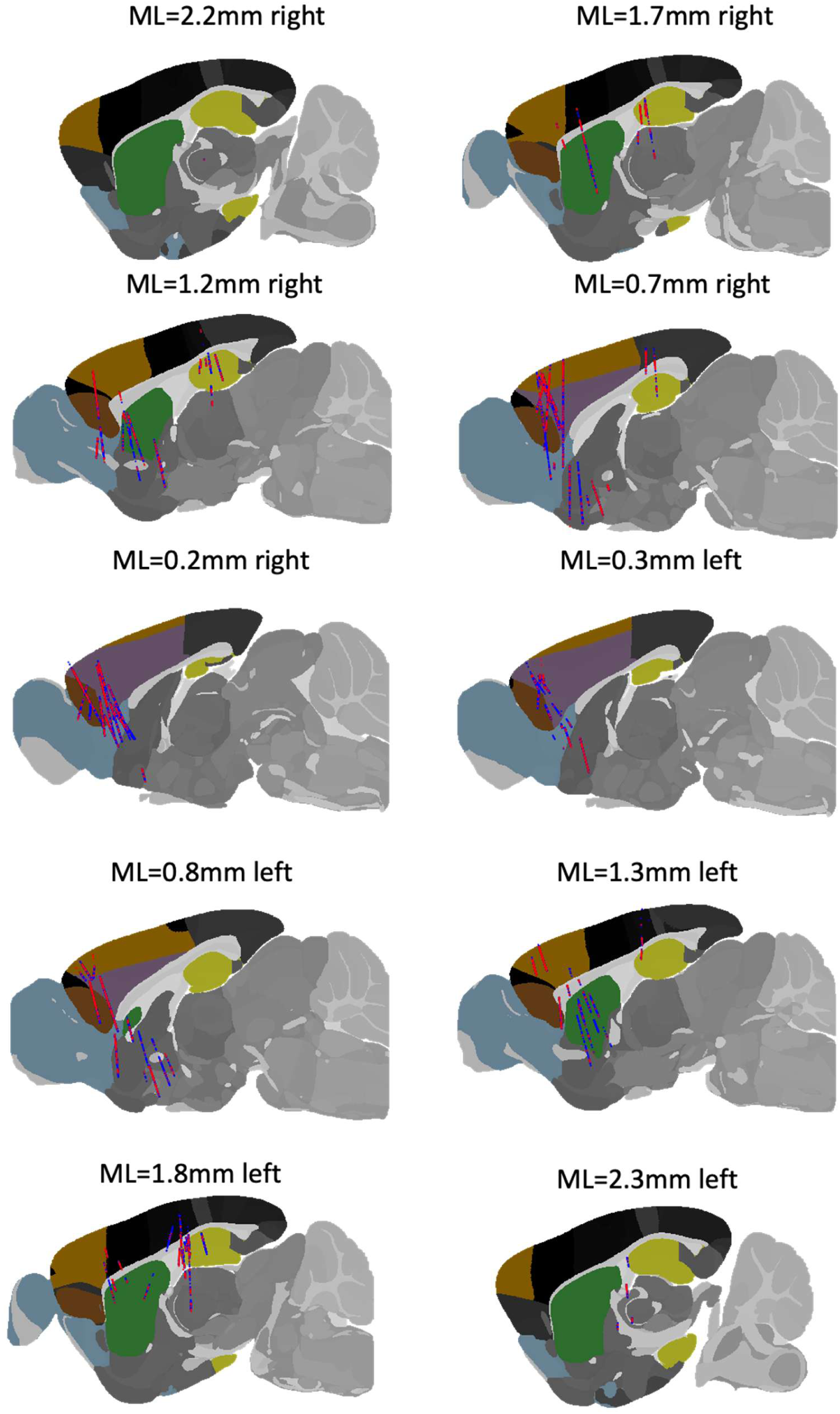
Spatial map of recording sites and choice correlation after the movement initiation. The maps show sagittal brain sections from the Allen Brain Atlas. The brain regions of interest are highlighted: MOs (orange), mPFC (lilac), orbital area (brown), dorsal hippocampus (yellow), dorso-medial striatum (green), olfactory areas (blue). Units that have significant (>0.15) correlation with choice after the movement has begun are marked red and the ones that do not are marked dark blue.

**Supplementary figure 9.**
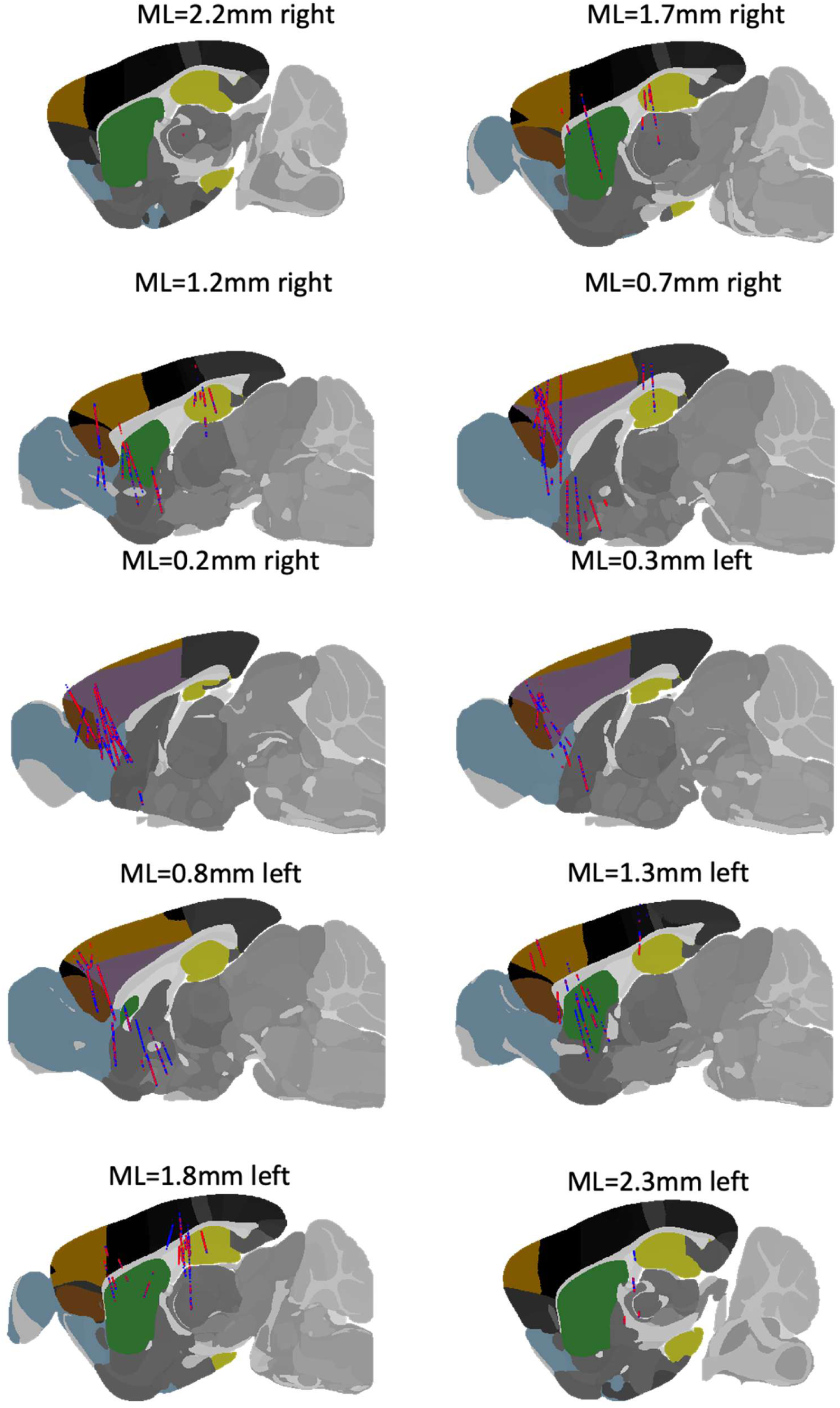
Spatial map of recording sites and feedback correlation after the feedback delivery. The maps show sagittal brain sections from the Allen Brain Atlas. The brain regions of interest are highlighted: MOs (orange), mPFC (lilac), orbital area (brown), dorsal hippocampus (yellow), dorso-medial striatum (green), olfactory areas (blue). Units that have significant (>0.15) correlation with choice after the movement has begun are marked red and the ones that do not are marked dark blue.

**Supplementary figure 10.**
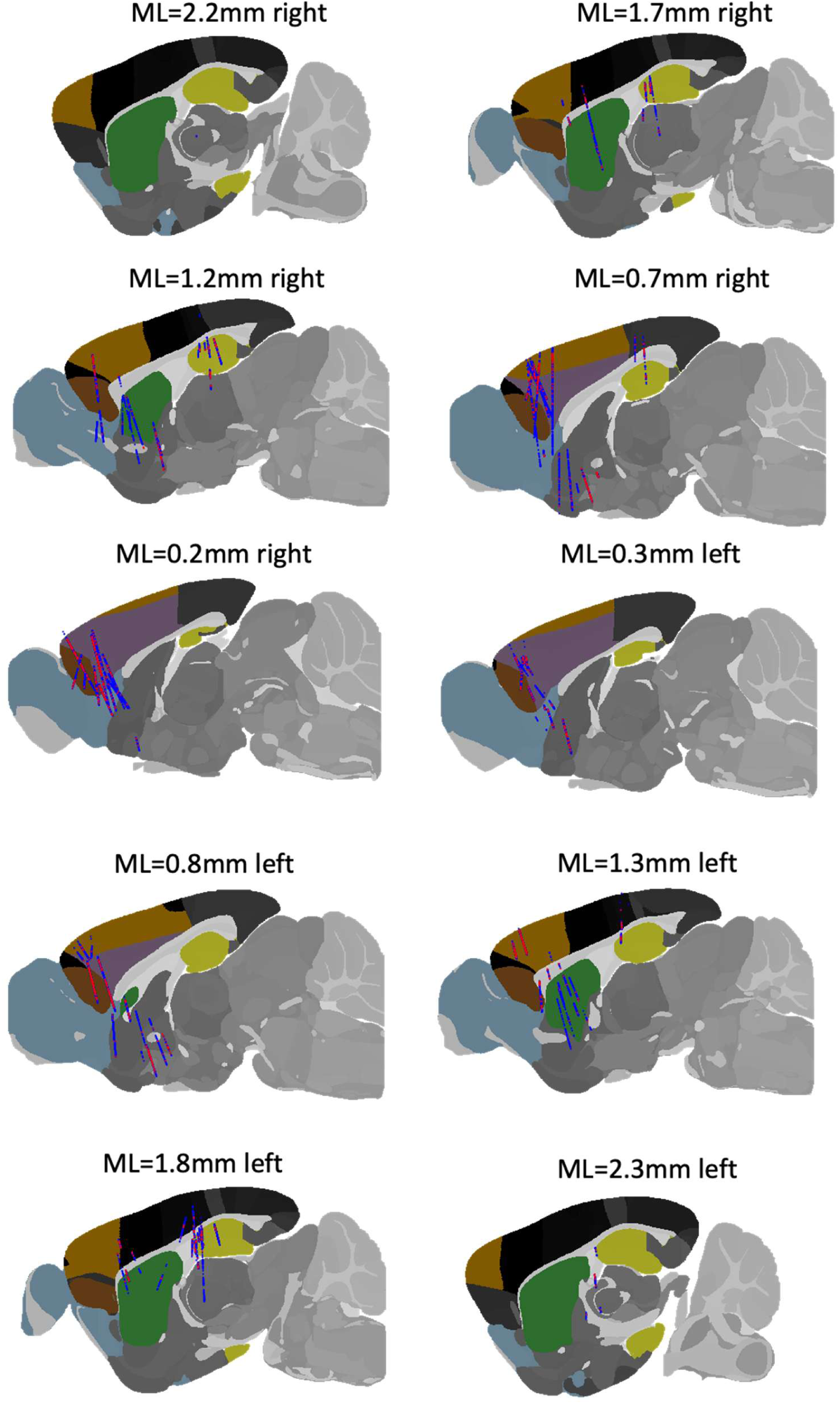
Spatial map of recording sites and choice correlation during the fixation period. The maps show sagittal brain sections from the Allen Brain Atlas. The brain regions of interest are highlighted: MOs (orange), mPFC (lilac), orbital area (brown), dorsal hippocampus (yellow), dorso-medial striatum (green), olfactory areas (blue). Units that have significant (>0.15) correlation with choice during the fixation period are marked red and the ones that do not are marked dark blue.

**Supplementary figure 11.**
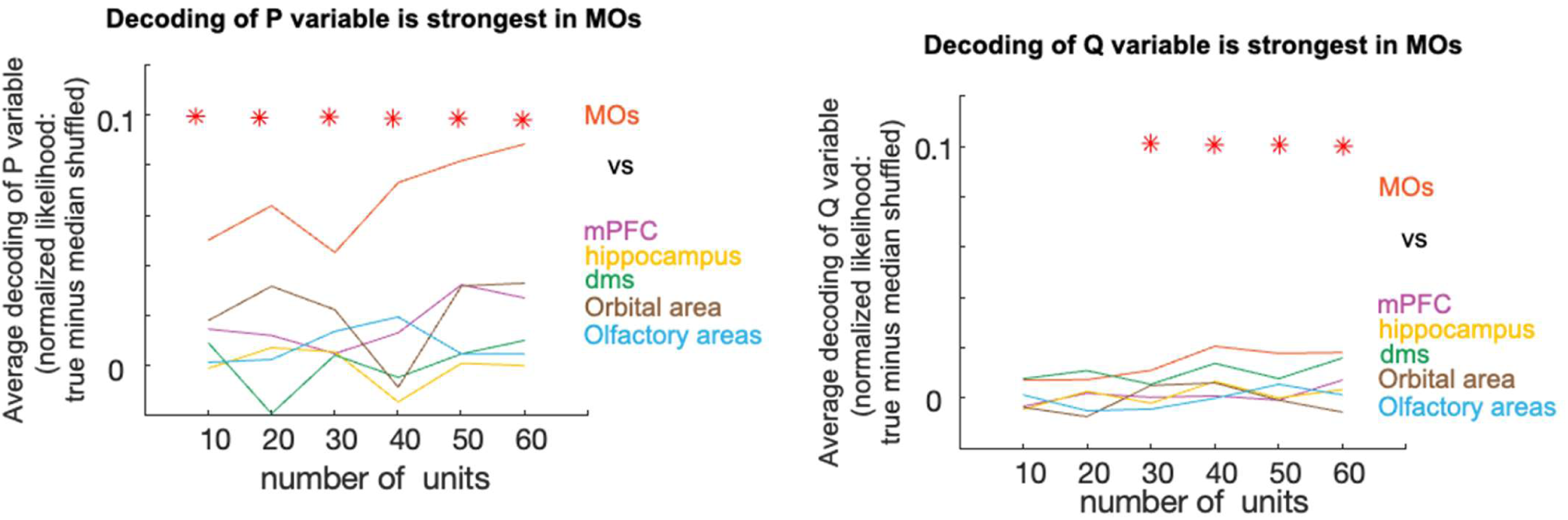
Decoding P and Q variables from fixation-period activity of equally-sized random subsamples of neurons from each region. X-axis: number of units in subsample; Y-axis, mean log-likelihood relative to permuted sessions. Red stars: MOs significantly different to other regions (p<.01; t-test).

**Supplementary figure 12.**
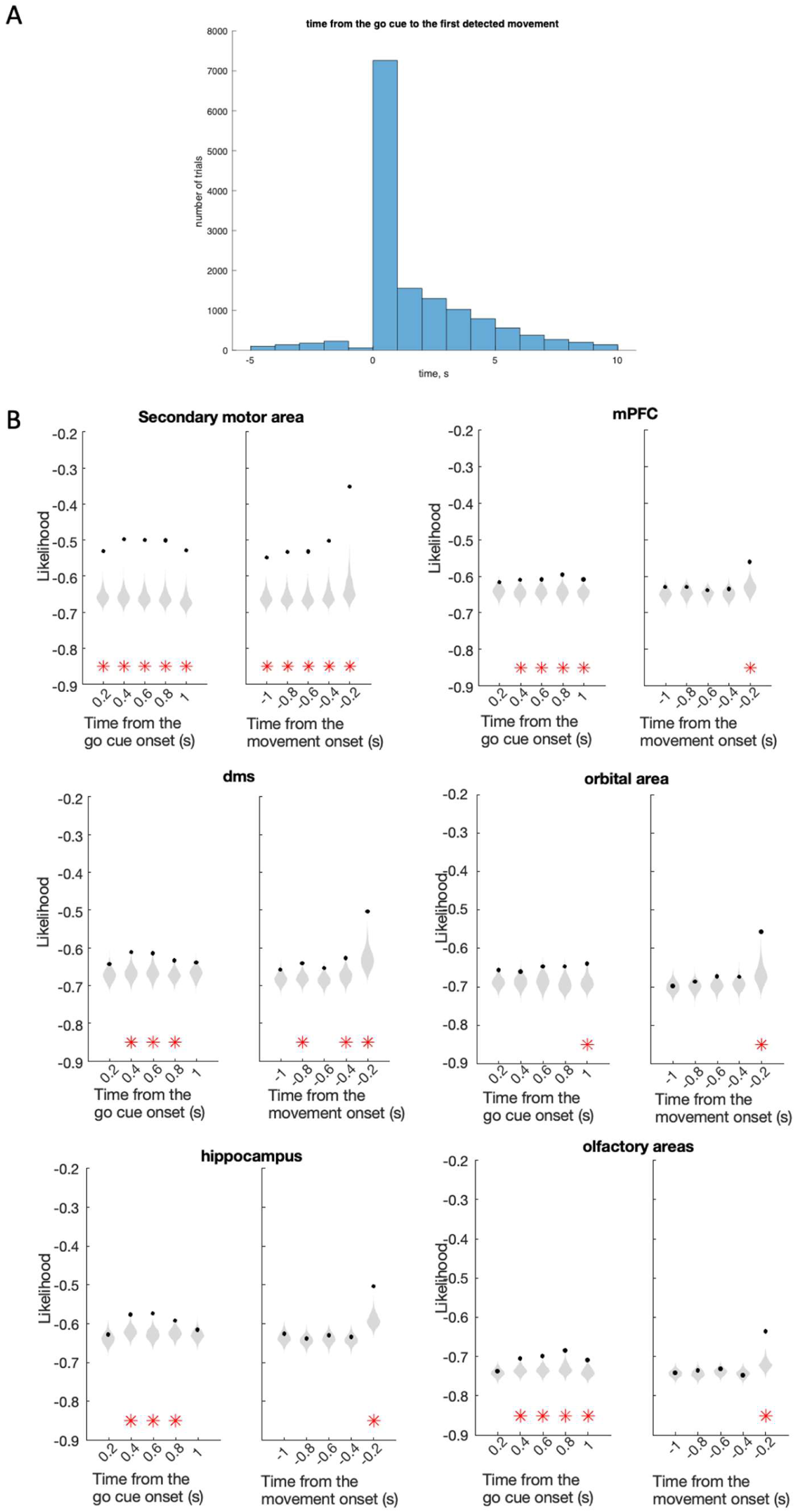
A, Bar graph showing the distribution of times between the first detected movement on a trial and the go cue time. B, Decoding of upcoming choice from population activity. Each column shows decoding of choice (logistic regression). For each region the decoding is done aligned to the go cue (left) and to the movement onset (right). X-axis: time epoch; 200 ms epochs after the go cue (left) or before movement onset (right). Black dots: mean prediction of actual value across sessions; gray violins, null distribution from session permutation. Red stars: p<0.01, session permutation test.

**Supplementary figure 13.**
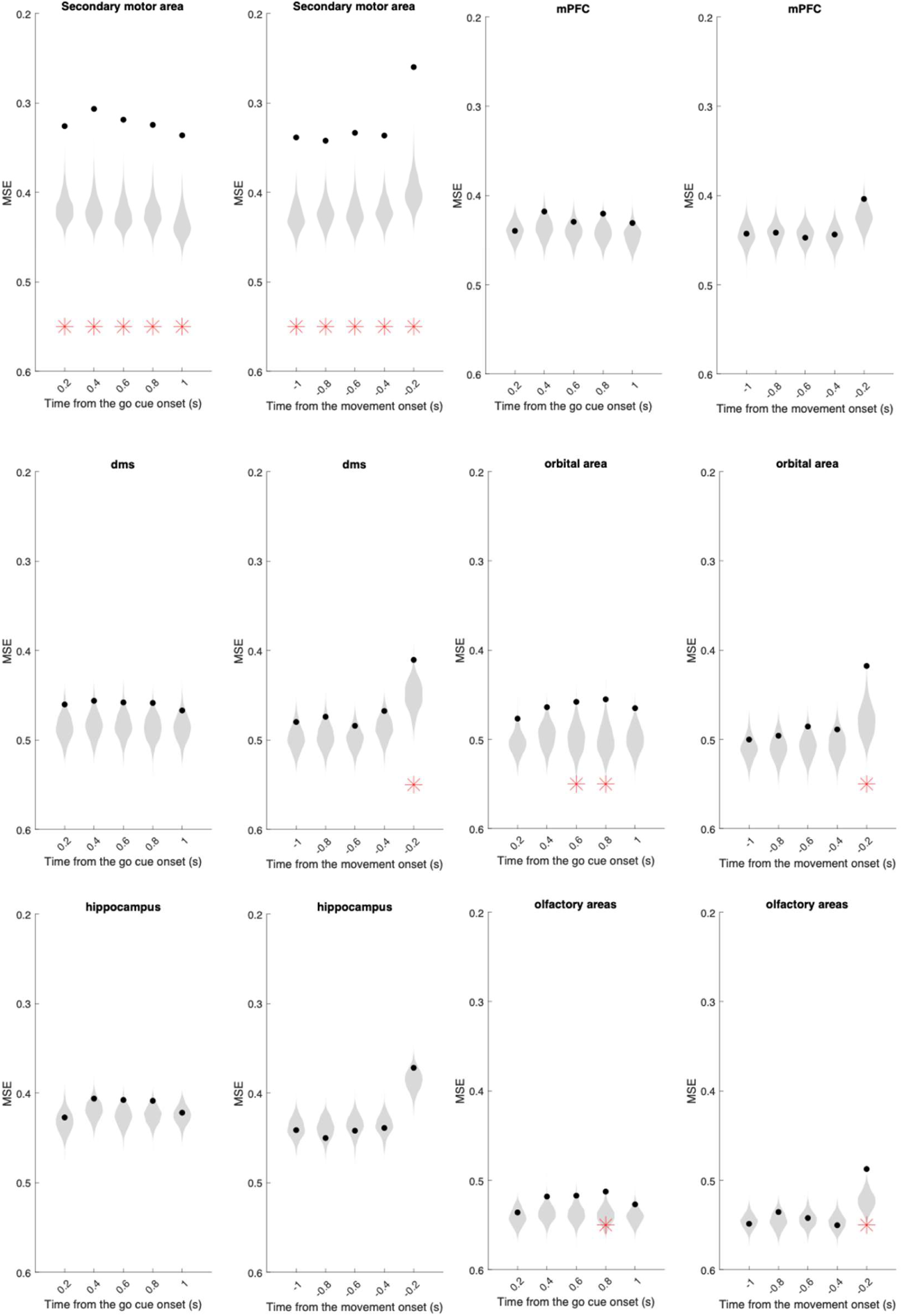
Decoding of P variable of the PR model from population activity. Each column shows decoding of P variable (linear regression). For each region the decoding is done aligned to the go cue (left) and to the movement onset (right). X-axis: time epoch; subsequent 200 ms epochs from go cue (left) or from movement onset (right). Black dots: mean prediction of actual value across sessions; gray violins, null distribution from session permutation. Red stars: p<0.01, session permutation test.

**Supplementary figure 14.**
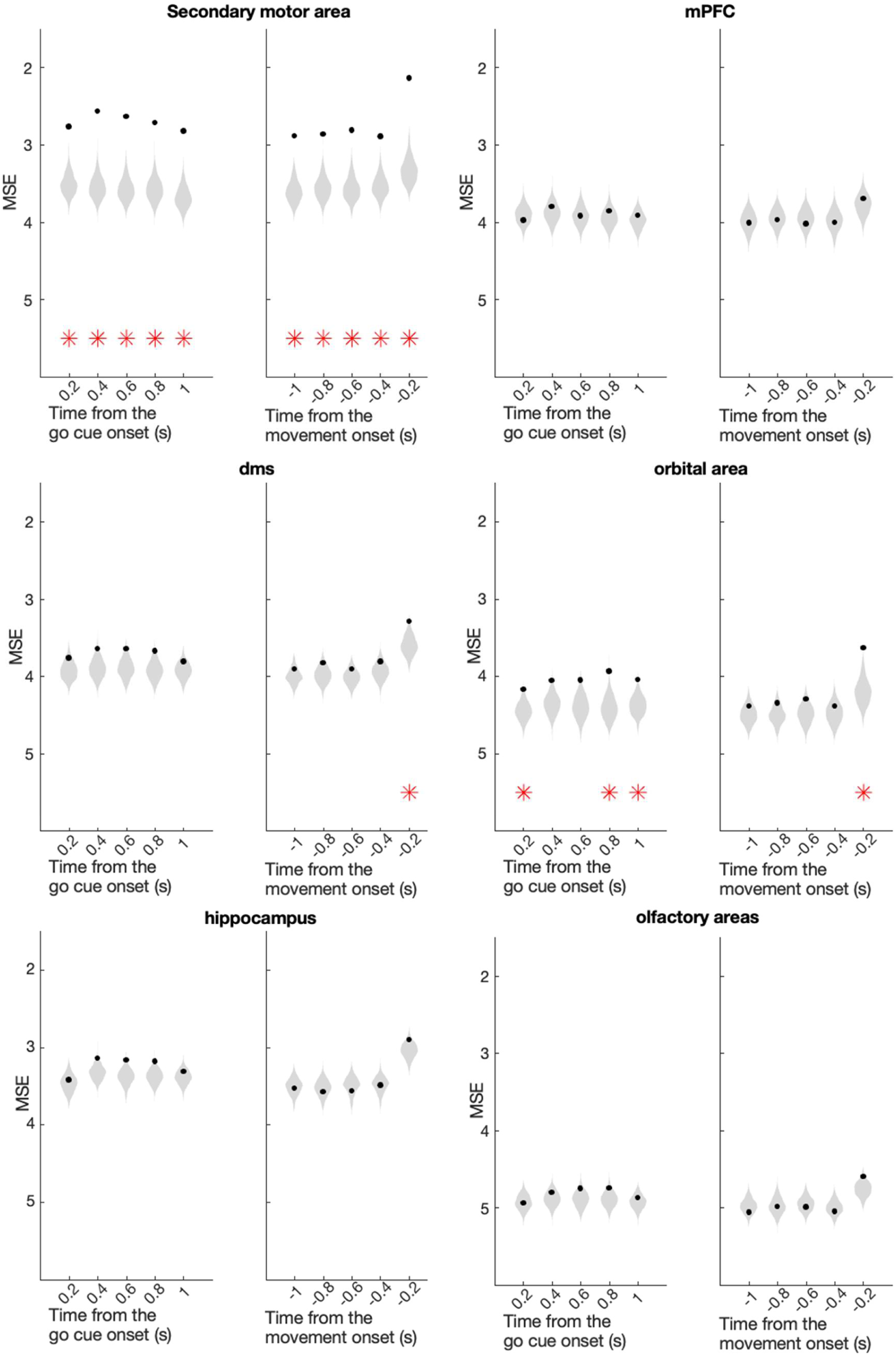
Decoding of P+R variable of the PR model from population activity. Each column shows decoding of P+R variable (linear regression). For each region the decoding is done aligned to the go cue (left) and to the movement onset (right). X-axis: time epoch; subsequent 200 ms epochs from go cue (left) or from movement onset (right). Black dots: mean prediction of actual value across sessions; gray violins, null distribution from session permutation. Red stars: p<0.01, session permutation test.

**Supplementary figure 15.**
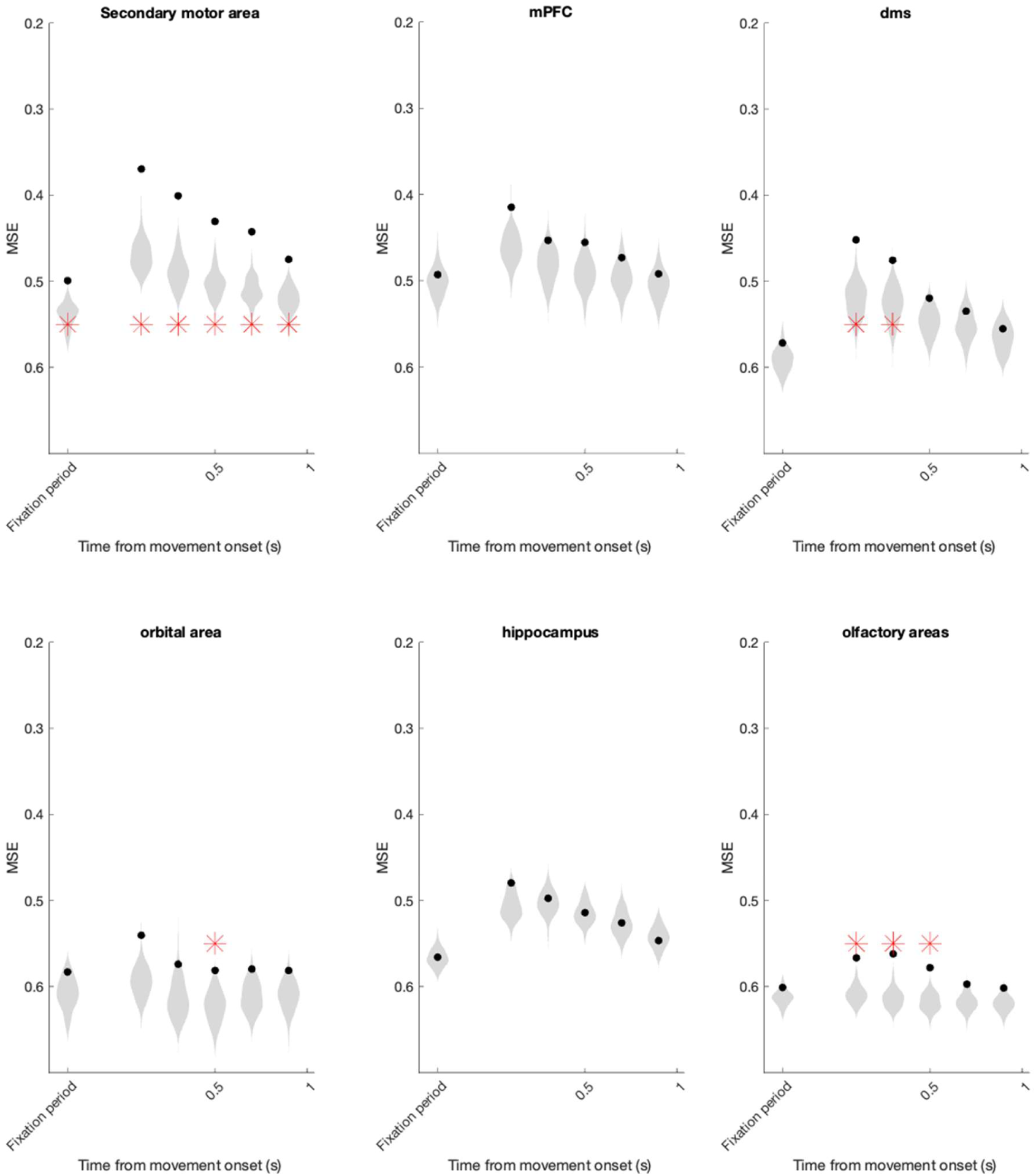
Decoding of perseveration variable of the PR model from population activity using LASSO regression. X-axis: time epoch; fixation period (0-200 ms before Go Cue), and subsequent 200 ms epochs from movement onset. Black dots: mean prediction of actual value across sessions; gray violins, null distribution from session permutation. Red stars: p<0.01, session permutation test.

**Supplementary figure 16.**
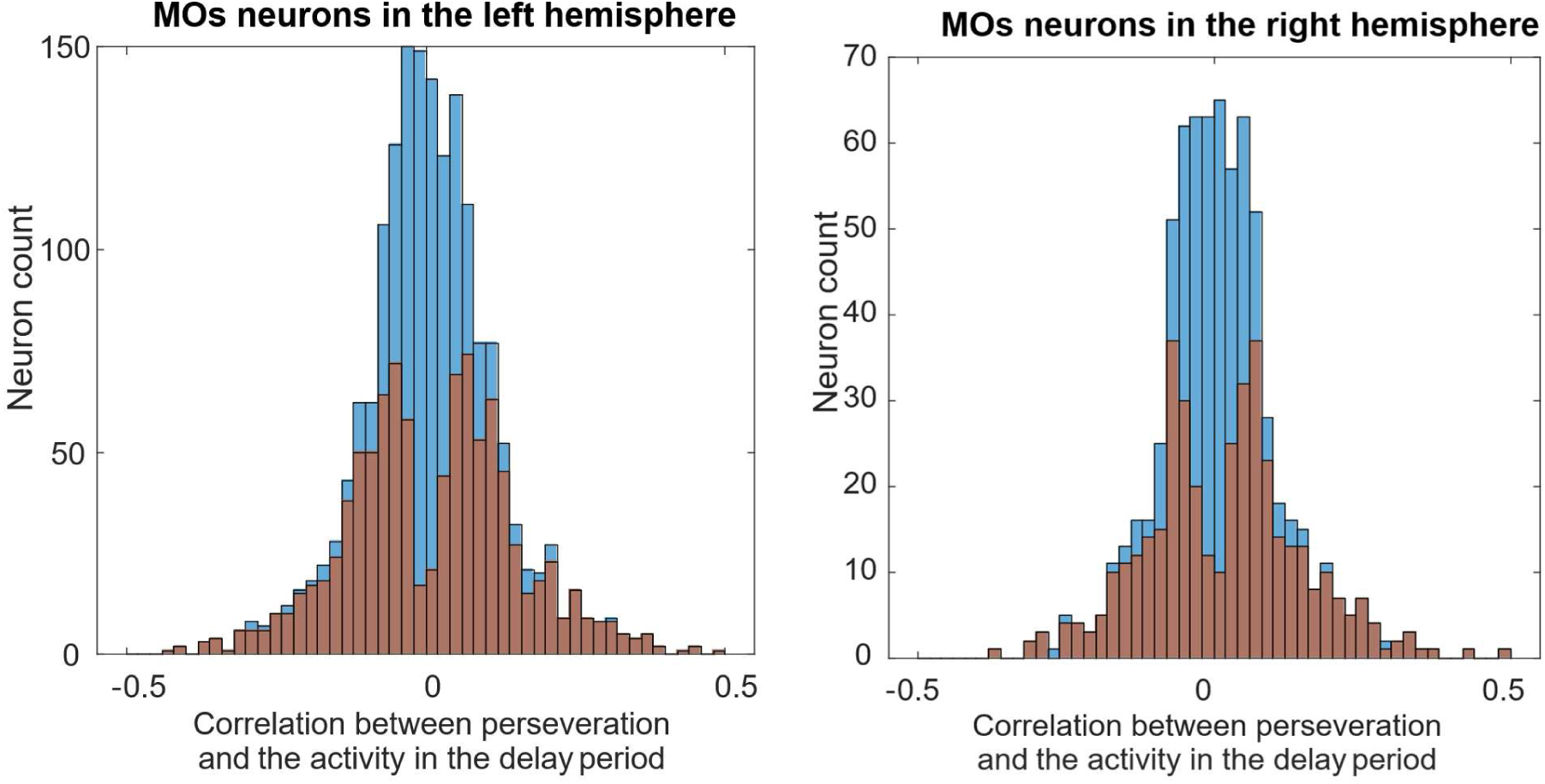
Left, correlation between activity of MOs clusters in the left hemisphere during the delay period and the perseveration variable. Blue, all units; orange, significant units according to session permutation test. Right, same for MOs units in the right hemisphere.

**Supplementary figure 17.**
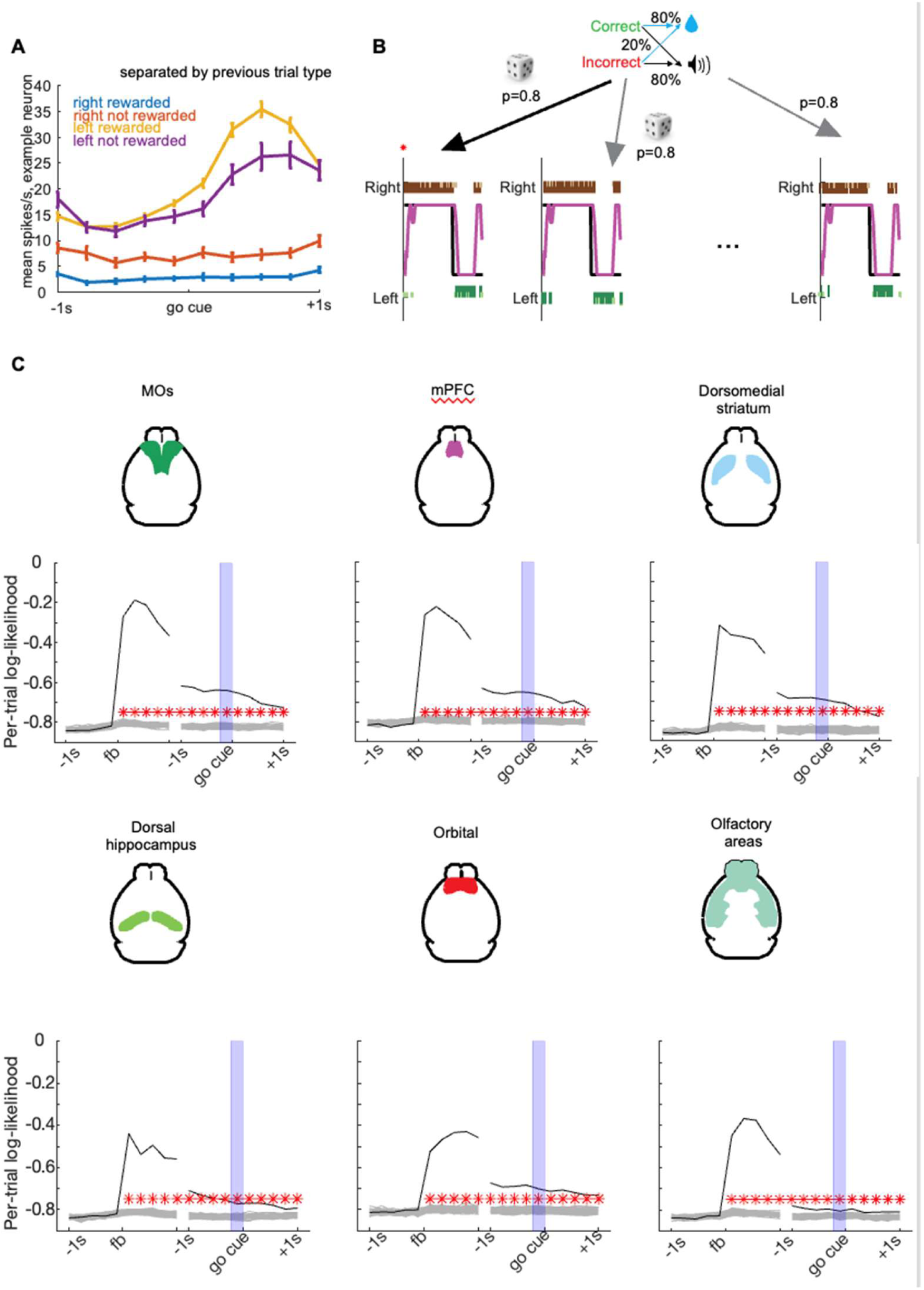
Causal effect of outcome (i.e. reward or white noise) on neural activity. **A**, Activity of an example neuron depends on the outcome of the previous trial. **B,** To show a causal effect we took advantage of the fact that feedback delivery was randomized (80% or 20% reward probabilities for correct or incorrect choices). For each trial, we generated a null ensemble by resampling the outcome at random, with the same probabilities as in the original task, given animals’ choices and correct responses. **C,** Accuracy of outcome prediction for the real session and the null ensemble. Black line shows log likelihood for predicting the actual outcome from neural population activity in a 200 ms time bin aligned to outcome time (left) or to the subsequent trial’s go cue (right). Blue shading indicates the fixation period. Gray lines indicate null ensemble predicting randomly regenerated outcome sequences, drawn using the same behavior-dependent probabilities as the actual sequence. Red stars indicate time bins in which the likelihood for predicting the actual choice exceeds 99% of the null ensemble. Note that even if the neural activity correlates with a prediction of upcoming reward magnitude, this analysis could not show a causal effect of outcome on activity before outcome delivery: because the null ensemble was randomly generated with the same probabilities as the actual outcome, there is no way that neural activity could predict the actual outcome better than null, even if it could predict whether the animal would choose the correct or incorrect side.

**Supplementary figure 18.**
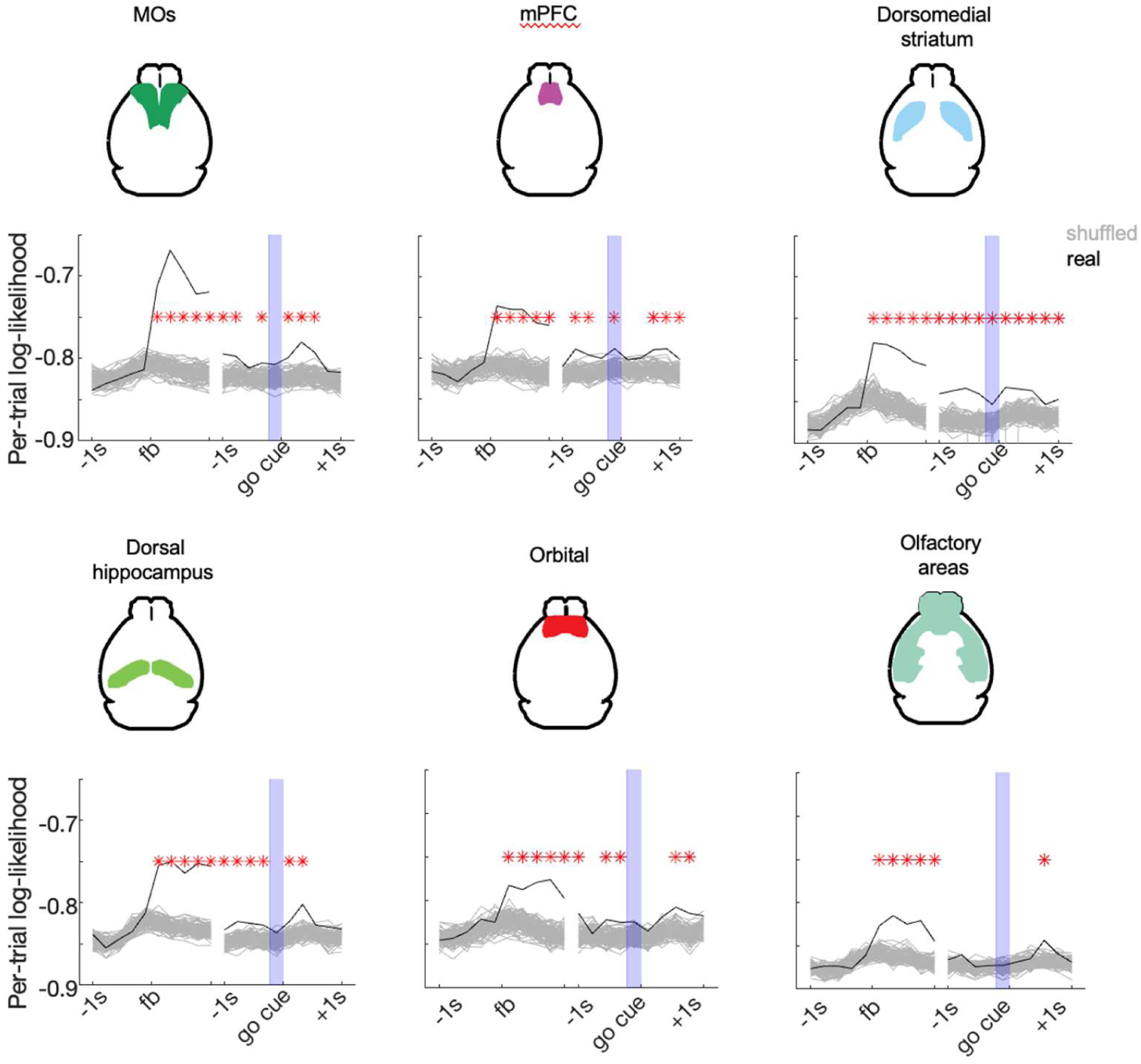
Causal effect of choice-feedback interaction on neural population activity, assessed by the same method as Figure 6. The analysis proceeds exactly as in Supp Fig. 6, but predicting the choice-feedback interaction rather than just the outcome value. Again, black lines indicate prediction of actual value, gray lines indicate the null ensemble obtained by resampling outcome values with the probabilities determined by the mouses actual choices, then recomputing the choice-outcome interaction. Blue rectangle, fixation period.

**Supplementary figure 19.**
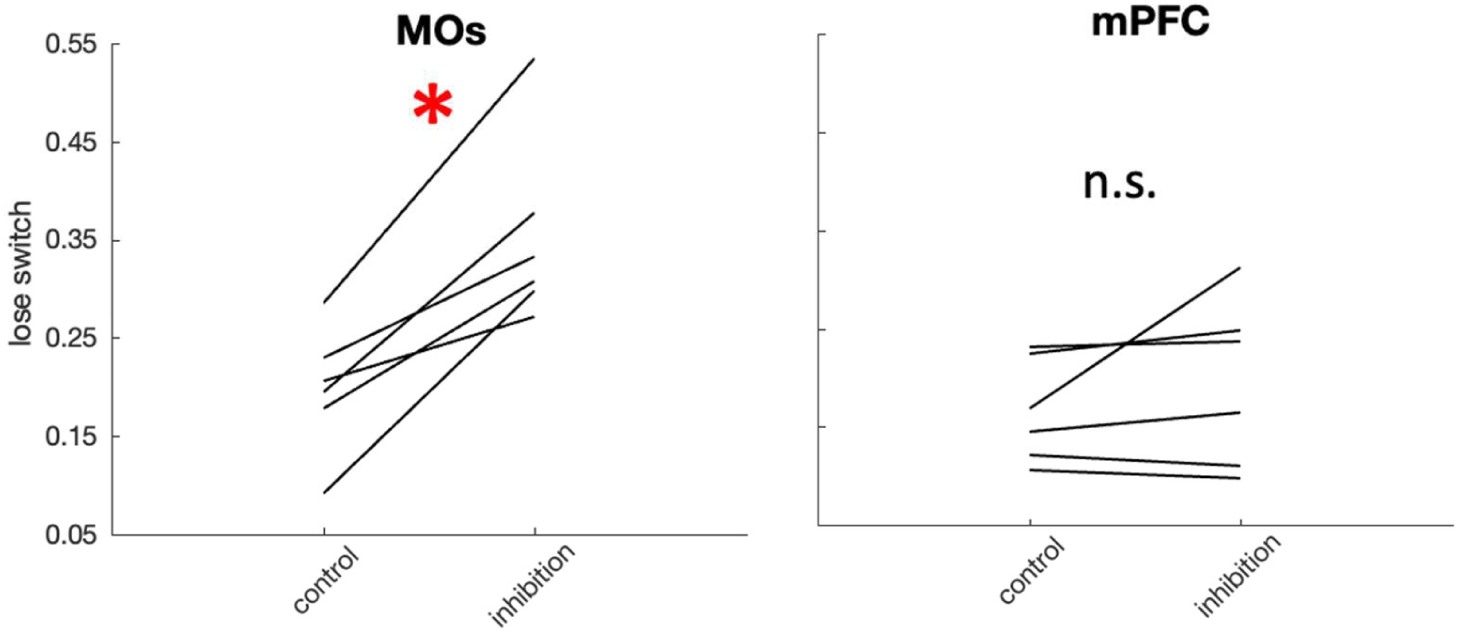
Probability of switching choice after no reward in no laser (control) and laser (inhibition) condition. Laser stimulation delivered upon the go cue.

**Supplementary figure 20.**
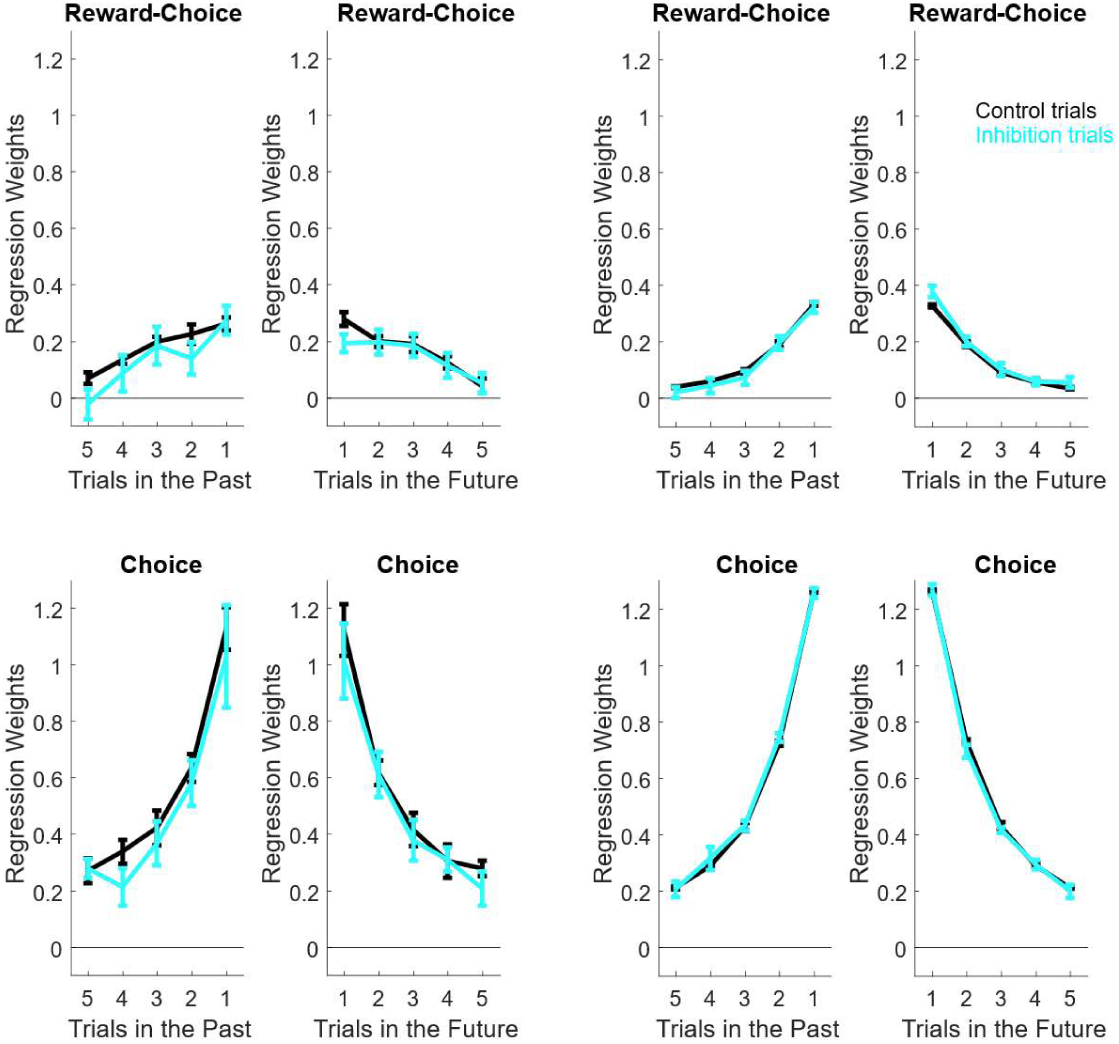
mPFC activity is not required for perseveration. Left 4 panels, Trial history regression weights for predicting mouse choice using past choices (choice weights), and choice-reward interaction (reward-choice weights), for laser (cyan) and no laser (black) conditions. Left column shows weights from trials N-n to trial N, where inactivation was delivered to mPFC on trial N; right column shows weights from trial N to trial N+n, where inactivation was delivered to mPFC on trial N. Right 4 panels show the same but for simulated data from PR model with no effect.

**Supplementary figure 21.**
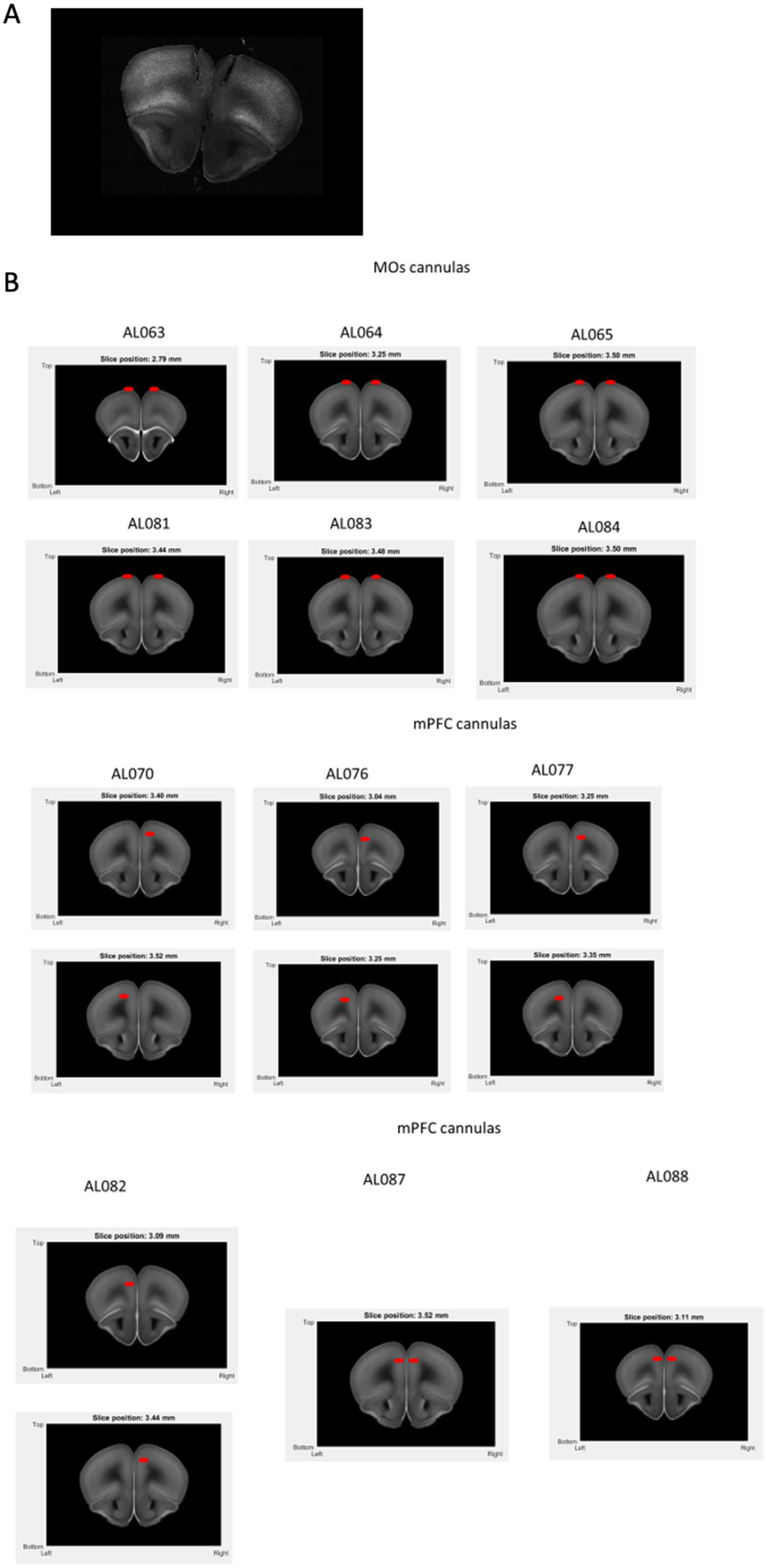
**A**, Example slice containing two cannula tracks (animal AL088). **B**, Estimated positions of the bottom of each cannula for optogenetic experiments (see methods).

**Supplementary figure 22.**
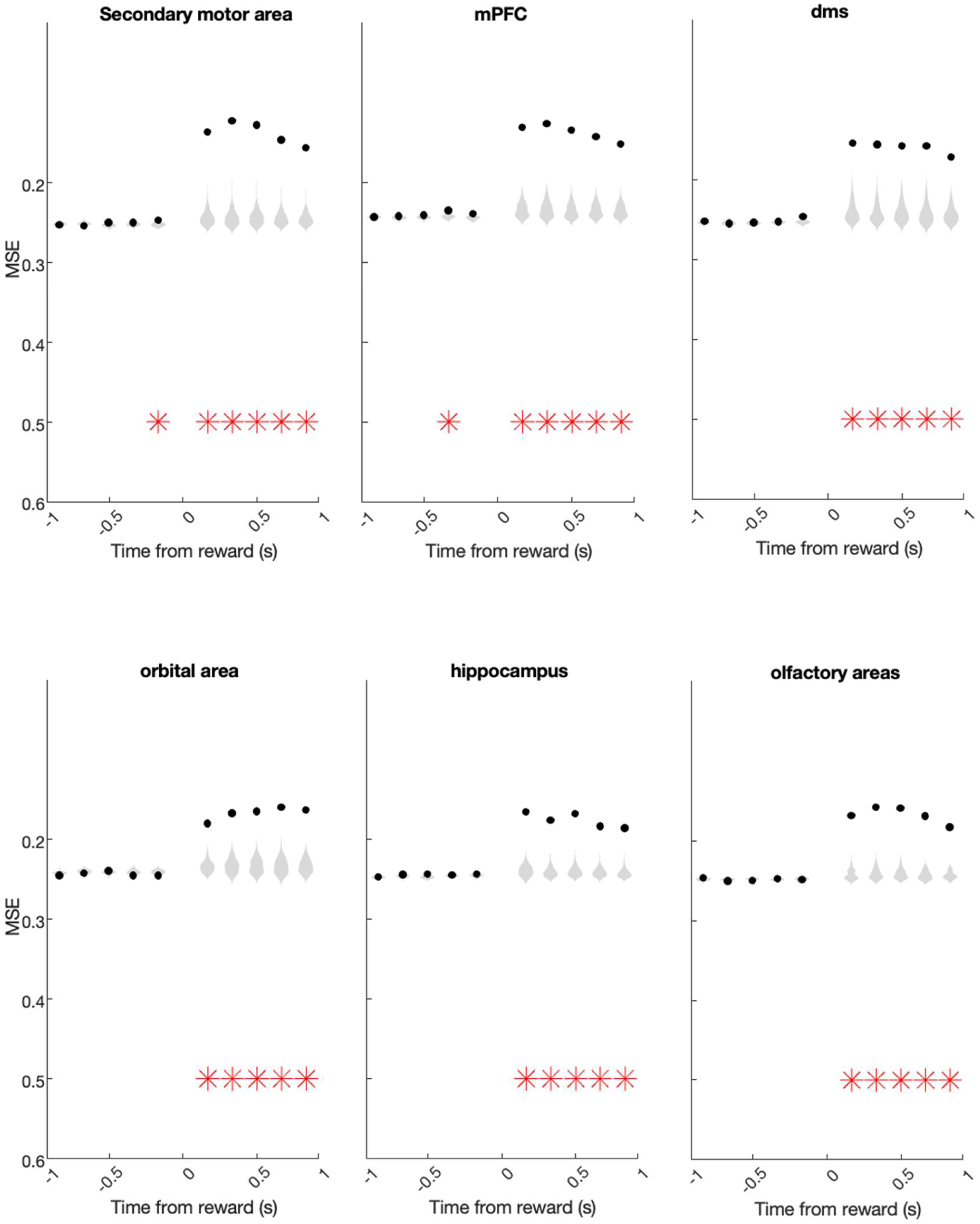
Decoding of reward prediction error (RPE), defined as actual reward minus the prediction of the standard Q-learning model (Methods) from population activity. Each plot shows decoding of RPE (linear regression). For each region the decoding is done aligned to the reward time. X-axis: time epoch; subsequent 200 ms epochs from reward time. Black dots: mean prediction of actual value across sessions; gray violins, null distribution from session permutation. Red stars: p<0.01, session permutation test.

## Notes

### Competing Interest Statement

The authors have declared no competing interest.

### Summary of Updates

Additional supplementary analyses to clarify concerns raised during peer review; expanded discussion of interpretation and previous literature.

